# Embracing Uncertainty Reshapes the ETHYLENE INSENSITIVE2-Activated Ethylene Signaling Framework

**DOI:** 10.1101/2024.06.13.598799

**Authors:** Hangwei Zhao, Ying Zhang, Yuying Chen, Chenrunshu Wang, Qian Liu, Jingyi Zhang, Chi-Kuang Wen

## Abstract

Signal transduction of the gaseous plant hormone ethylene by ETHYLENE INSENSITIVE2 (EIN2) is proposed to be regulated at multiple levels. EIN2 is phosphorylated by CONSTITUTIVE TRIPLE-RESPONSE1 (CTR1), subject to the F-Box proteins [EIN2-TARGETING PROTEIN1 (ETP1) and ETP2]-mediated degradation, and cannot activate ethylene signaling. Ethylene prevents EIN2 phosphorylation and degradation, and EIN2 accumulates and activates the signaling. MAOHUZI3 LIKE1 (MHL1) and MHL2 stabilize EIN2, and the *mhl1 mhl2* double mutation confers ethylene insensitivity. Here, we reported that the regulation of EIN2-activated ethylene signaling can be independent of CTR1-mediated phosphorylation, ETP1/ETP2-directed degradation, and MHL1/MHL2-dependent stabilization. Both the *etp1 etp2* double mutant and wild type showed identical ethylene dose-response curves, and the nontreated *mhl1 mhl2* double mutant displayed substantial growth inhibition. The reported ethylene-insensitive root phenotype of *mhl1 mhl2* seedlings requires sucrose and is associated with unknown loci. The ethylene receptor ETHYLENE RESPONSE1 (ETR1) and EIN2 interact at the rough endoplasmic reticulum. We propose that the docking of CTR1 to ETR1 promotes receptor signaling, which inhibits EIN2-activated ethylene signaling. The discrepancy between our findings and the current model is discussed. Our findings may disrupt the knowledge boundary of the present molecular model, developing a niche for findings outside the framework to advance our understanding of ethylene signaling.

## Introduction

The signal transduction of the plant hormone ethylene has been investigated for approximately three decades, with the mechanism of the linear genetic pathway elucidated through molecular biology, cell biology, and omics studies. Although the underlying biochemical properties are poorly understood, ethylene signaling is proposed to involve interlaced multilevel regulation, including protein phosphorylation, proteolytic cleavage-coupled protein transport, translation, and protein stability (3–7). The five ethylene receptor isoforms are classified into two subfamilies: ETHYLENE RESPONSE1 (ETR1) and ETHYLENE RESPONSE SENSOR1 (ERS1) are subfamily I members, and ETR2, ETHYLENE INSENSITIVE4 (EIN4) and ERS2 are subfamily II members (8, 9). The five isoforms activate the Raf-like protein CONSTITUTIVE TRIPLE-RESPONSE1 (CTR1) in the absence of ethylene (10, 11), and CTR1 phosphorylates the endoplasmic reticulum (ER)-associated EIN2, which contains the natural resistance-associated macrophage protein (Nramp)-like transmembrane domain at the amino terminus (3, 4). Phosphorylated EIN2 is thought to remain at the ER and undergo degradation mediated by the F-box proteins EIN2-TARGETING PROTEIN1 (ETP1) and ETP2 (2–4). Ethylene treatment prevents ethylene receptor signal output and CTR1 activation, allowing nonphosphorylated EIN2 to be stabilized by MAOHUZI3-LIKE1 (MHL1) and MHL2 and to be proteolytically cleaved at the Ser645 residue to release and transport the carboxyl portion (EIN2-C or EIN2^646-1294^) to the nucleus (3, 5, 13). EIN3-BINDING F-BOX1 (EBF1) and EBF2 are F-box proteins that mediate the degradation of the master transcription factors EIN3 and EIN3-LIKE1 (EIL1) (14–16). EIN2-C targets the 3’-untranslated region (3’-UTR) of the *EBF1* and *EBF2* mRNAs, and the EIN2-*EBF1/EBF2* ribonucleoprotein complex integrates into the processing body (PB), presumably via liquid-liquid phase separation. *EBF1*/*EBF2* mRNA translation is repressed (6, 7), and EIN3/EIL1 accumulates and induces ethylene response genes.

CTR1-mediated EIN2 phosphorylation was determined by phosphoproteomic and *in vitro* phosphorylation studies (3, 4). The serine residues at positions 645, 739, 743, 744, 747, 757, 1283, and 923/924 and the threonine^742^ residue are phosphorylated. Among these residues, S645 and S924 are conserved across several plant species, with high and moderate degrees of confidence in position assignment, respectively, and S1283 has the least amount of conservation (4). An *in vitro* assay revealed substantially reduced (by >85%) EIN2^479-1294^ phosphorylation when both the Ser645 and Ser924 residues were replaced with alanine, indicative of differential or cooperative phosphorylation of the other residues. Mass spectrometry analysis of the *in vitro* kinase reaction identified six EIN2 phosphopeptides, of which the S659 and T819 residues were not identified via phosphoproteomics. These studies indicate that at least four sites (S645, S757, S924, and S1283) are phosphorylated *in vivo* by CTR1 (3).

Qiao *et al.* investigated the biological significance of CTR1-mediated phosphorylation and proposed the present framework of EIN2-activated ethylene signaling (5). While this model perfectly describes the cytoplasmic-nuclear transport of EIN2 as a carrier of the ethylene signal, there are obvious caveats to the study and contradictions accumulated along with follow-up studies. A single mutation of the Ser645 phosphorylation residue is sufficient to determine ethylene signaling, and expression of the *EIN2^S645A^-GFP*-*HA* transgene confers constitutive ethylene signaling. Western blots show that the nonphosphorylated EIN2^S465A^ protein is predominantly uncleaved, and GFP fluorescence is localized to the nucleus. The association between the amount of full-length EIN2^S645A^ and nuclear fluorescence implies that full-length EIN2^S645A^, but not cleaved EIN2-C, activates ethylene signaling in the nucleus. Moreover, if EIN2 is cleaved at Ser645, the Ser645Ala replacement conceivably eliminates the cleavage site, prevents the cleavage, and nullifies EIN2 functionality (17). The activity of EIN2^S645A^ and the absence of the semi-tryptic EIN2^630-645^ peptide do not support the EIN2 cleavage and transport model (17, 18). In addition to these caveats, the results from independent studies are inconsistent with those of Qiao *et al*. (5). Resulting from ethylene-induced EIN2 stabilization, Qiao *et al*. reported an elevation of EIN2 levels, (2), which contradicts the finding that ethylene reduces EIN2 levels, resulting from EIN2 cleavage and nuclear transport (3, 5). In contrast to the single cleavage-coupled nuclear transport model, two independent follow-up studies reported the dual localization of full-length EIN2 in the nucleus and cytoplasm, and EIN2 can be processed into highly complex fragments in an ethylene-independent manner (18, 19). The cleavage complexity is also determined for the rice homolog OsEIN2 and the endogenous Arabidopsis EIN2 (13, 20). In contrast to reports that the hypothetical phosphomimicking EIN2^S645D^ or EIN2^S645E^ fails to activate ethylene signaling (5), Zhang *et al.* showed that the expression of the wild-type *EIN2*, *EIN2^S645A^*, *EIN2^S645D^ ^S924D^*, or *EIN2^S645E^ ^S924E^*transgene complements the *ein2* mutation and confers various degrees of constitutive ethylene signaling. EIN2 or these variants are localized to the cytoplasm and nucleus independent of ethylene treatment (18), supporting the idea that EIN2 functionality can be independent of the Ser645/Ser924 residues. The degree of constitutive ethylene signaling is associated with EIN2 levels but not with the phosphorylation residues (3, 18).

In addition to these contradictory reports, the present model does not explain CTR1-independent ethylene signaling, where the ETR1 N-terminus ETR1^1-349^ reverses the *ctr1* mutant phenotype in a *REVERSION-TO-ETHYLENE SENSITIVITY1* (*RTE1*)-dependent manner (21), indicating other possibilities for ethylene signal transduction. This finding aligns with the extreme *etr1 ers1* mutant phenotype in contrast to the typical *ctr1-3* constitutive triple-response phenotype (9, 11). In this study, we provided experimental evidence that the four CTR1-mediated phosphorylation residues are nonessential for EIN2 functionality, that the ethylene receptor ETR1 can prevent EIN2-activated ethylene signaling independent of CTR1, and that ethylene signaling activation is independent of the regulation of the F-box proteins ETP1 and ETP2 or the membrane proteins MHL1 and MHL2. We discuss the discrepancy between different studies, by which findings hidden under Occam’s broom can be unveiled and a niche can be developed to expand the knowledge boundary of ethylene signaling. A molecular model is proposed and discussed to explain ethylene signaling.

## Results

### Ethylene Signaling Is Independent of CTR1-mediated EIN2 Phosphorylation

In contrast to reports that the non-phosphorylatable S645A or S645A S924A but not the phosphomimic Ser645D/E mutation of EIN2 leads to constitutive ethylene signaling (3, 5), we previously reported that the *EIN2*, *EIN2^S645A^*, *EIN2^S645A^ ^S924A^*, or *EIN2^S645D/E^ ^S924D/E^* transgene complemented the *ein2* loss-of-function mutation, conferring various degrees of constitutive ethylene signaling, and these transgenic plants were ethylene responsive (18). This finding implies that the role of EIN2 phosphorylation in ethylene signaling activation is unclear (3, 5).

Among the many phosphorylation residues, CTR1 phosphorylates at least four serine residues of EIN2 (3, 4). To test the biological significance of CTR1-mediated phosphorylation in EIN2 functionality, the four phosphorylation residues were either replaced with the non-phosphorylated A (EIN2^quadrupleA^, designated EIN2^qA^), phosphomimic D/E (EIN2^qD^/EIN2^qE^), or removed (EIN2^qΔS^) (Fig. 1A). When expressed in the *ein2^W308*^* mutant, resulting from early termination at the eighth transmembrane helix of the Nramp domain, each transgene complemented the *ein2^W308*^* mutation and conferred varying degrees of constitutive ethylene signaling, as evidenced by the seedling hypocotyl growth inhibition phenotype. Ethylene treatment inhibited the seedling hypocotyl growth of these transgenic plants to a similar extent, suggesting the activation of ethylene signaling. As a control, the *EIN2-eYFP* transgene had a similar effect on *ein2^W308*^* seedling growth and ethylene responses (Fig. 1B and 1C). In line with the reported complexity of EIN2 cleavage (13, 18–20), our western blots revealed a similar cleavage complexity for EIN2 and those EIN2 variants, involving rabbit polyclonal antibodies or a mouse monoclonal antibody (Fig. 1D). These consistent results for the two types of antibodies exclude the possibility that EIN2 complexity arises from nonspecific epitope recognition. Although D or E replacement is hypothetically phosphomimicking (3, 5), it has yet to be determined whether these replacements realistically mimic or prevent phosphorylation. The minimal impact of the deletion or mutation of the four serine residues on ethylene responses suggests that the EIN2 variants that cannot be modified by CTR1 are not autonomously active, as proposed, and can be activated by ethylene.

**Figure 1.**
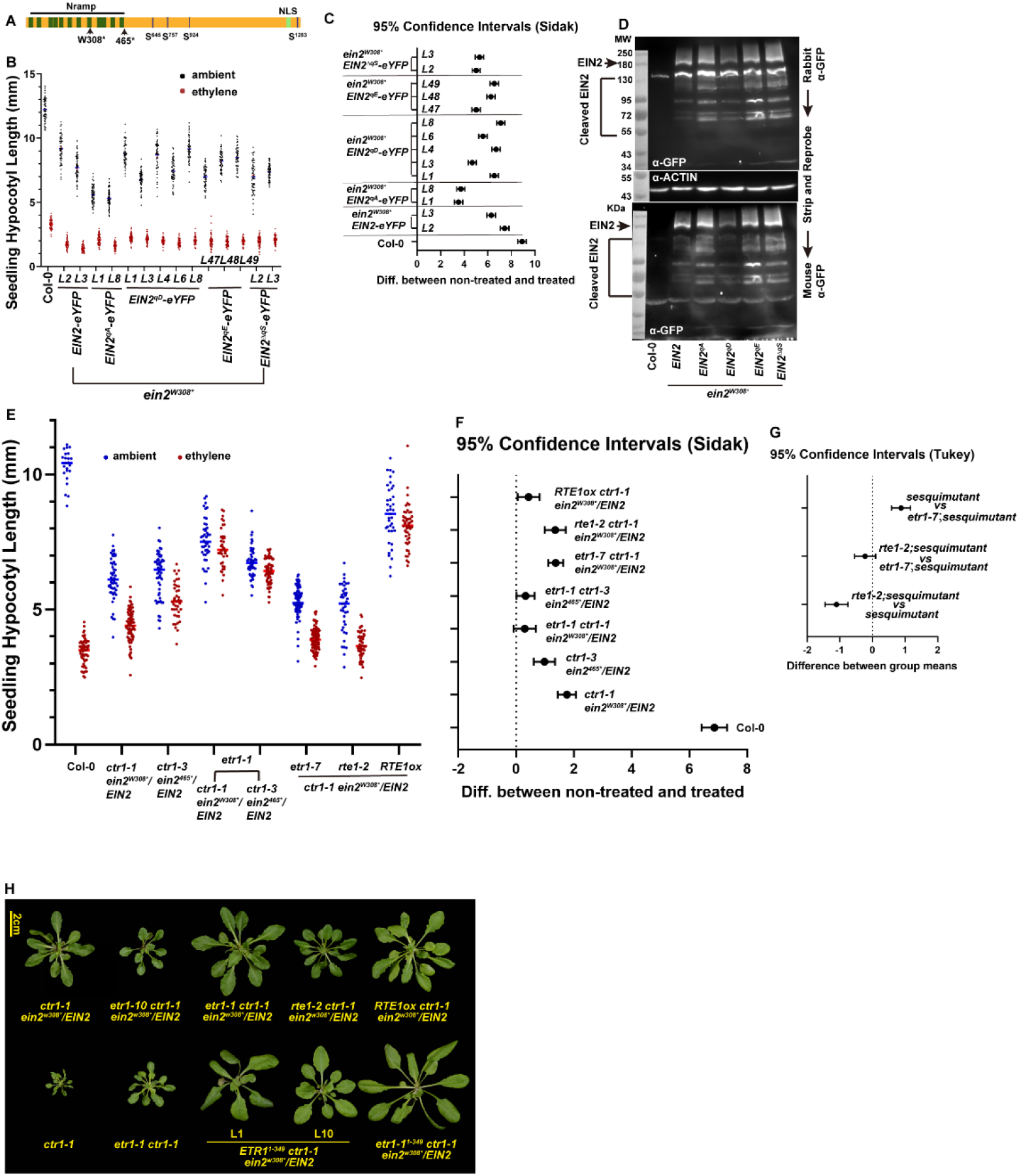
EIN2 activated ethylene signaling independent of phosphorylation. (A) An illustration of the EIN2 protein structure; the Nramp-like transmembrane domain, the four CTR1-mediated phosphorylation residues, and the nuclear localization signal (NLS) are indicated. The sites for the *ein2^W308*^*and *ein2^465*^* early terminations are denoted. Seedling hypocotyl measurements (*n*≥30) of the transgenic lines expressing the indicated *EIN2* transgenes in the *ein2^W308*^* mutant (B) and the hypocotyl mean differences (CI_0.95_) between the nontreated and ethylene-treated seedlings (C). The four serine residues were replaced with alanine (qA), aspartic acid (qD), glutamate (qE), or deleted (ΔqS). (D) Western blots showing the chemiluminescence of EIN2-eYFP and the eYFP-tagged EIN2 variants; the blot probed with the rabbit polyclonal anti-GFP antibodies were stripped and reprobed with the mouse monoclonal anti-GFP antibody. NS: nonspecific signals. ACTIN was used as the internal control. The original images can be found from Figure1D_source_data_1 and Figure1D_source_data_2. (E) Seedling hypocotyl measurements (*n*≥30) of the indicated genotypes revealed the ethylene responsiveness of the two *ctr1 ein2/EIN2* sesquimutants and the ethylene insensitivity conferred by the ethylene-insensitive *etr1-1* mutation or *RTE1* overexpression. (F) The mean hypocotyl differences (CI_0.95_) for the indicated genotypes between the nontreated and ethylene-treated seedlings. (G) The differences in the mean hypocotyl length (CI_0.95_) between the indicated genotypes indicate that *etr1-7* and *rte1-2* have similar effects on seedling growth. (H) The rosette phenotype of the sesquimutant and mutants with the indicated mutation or transgene. The data are presented as the means and standard errors (SEs), and the dots indicate individual measurements. The transgenes included the *EIN2* promoter for *EIN2* transgenes and the *ETR1* promoter for the *ETR1^1-349^*/*etr1-1^1-349^*transgene. CI_0.95_: 95% confidence interval.

In addition to the four CTR1-mediated phosphorylation residues, EIN2 can be differentially phosphorylated at other residues (4). Notably, the *ctr1-3* mutant is responsive to ethylene (18) and has a much weaker phenotype than the *etr1 ers1* mutant (8, 9), implying that EIN2 phosphorylation at residues independent of CTR1 plays a role in ethylene signaling. This other possibility is an alternative mechanism that prevents EIN2-activated ethylene signaling (9, 22–24). The latter scenario aligns with the prevention of ethylene-induced growth inhibition of the *ctr1-2* and *etr1-7 ctr1-2* mutants by the ethylene-insensitive *ethylene response1-1^1-349^*(*etr1-1^1-349^*) transgene (9, 21, 22), where the four CTR1-mediated phosphorylation residues of EIN2 are not modified. Notably, the artificially created ETR1-1^1-349^ fragment may not fully represent the ETR1 receptor, and whether EIN2 activity can be inhibited by ETR1 independent of CTR1 has yet to be investigated. The *ctr1-1 ein2^W308*^*/*EIN2* and *ctr1-3 ein2^465*^*/*EIN2* sesquimutants, of which the four serine residues of EIN2 are not phosphorylated in the absence of CTR1, exhibited a moderate degree of constitutive ethylene signaling and were responsive to ethylene (Fig. 1E and 1F) (18). The ethylene-insensitive *etr1-1* mutation strongly reversed the sesquimutant phenotype, and the *etr1-1 ctr1-1 ein2^W308*^*/*EIN2* and *etr1-1 ctr1-3 ein2^465*^*/*EIN2* mutants displayed much weaker seedling growth inhibition than the sesquimutants and were ethylene insensitive (Fig. 1E and 1F). Similarly, *REVERSION-TO-ETHYLENE SENSITYVITY1* (*RTE1*) overexpression, which promotes ETR1 receptor signaling and confers ethylene insensitivity, also prevents the ethylene response of the sesquimutant (Fig. 1E and 1F)(21, 22, 25). The *etr1-7* or *rte1-2* loss-of-function mutation enhanced the inhibition of seedling hypocotyl growth of the sesquimutant to a similar extent (Fig. 1E-1G; Tukey test). In line with the seedling mutant phenotype, *ETR1^1-349^*, *etr1-1^1-349^*, *etr1-1*, or *RTE1* overexpression substantially reversed the *ctr1-1 ein2^W308*^*/*EIN2* sesquimutant growth inhibition at the rosette stage, and the *etr1-7* or *rte1-2* mutation elevated the sesquimutant growth inhibition. Notably, the *etr1-1* mutation reversed *ctr1-1* mutant growth (Fig. 1H).

Our present data suggest that the four serine residues that are phosphorylated by CTR1 play little role in determining EIN2 activation, the activation of EIN2 and those variants requires ethylene, EIN2 proteolytic cleavage is highly complex and autonomous, and the ETR1 receptor can prevent EIN2-activated ethylene signaling in the absence of CTR1-mediated phosphorylation.

### ETR1 Mutations in the Histidine Kinase Domain Prevent EIN2-Activated Ethylene Signaling Independent of CTR1

In contrast to the reversion of the sesquimutant phenotype by *etr1-1* or *RTE1* overexpression, *etr1-1* marginally reverses the *ctr1-1* mutant phenotype, and *RTE1* overexpression has little effect on *ctr1-1* (21), which is indicative of *EIN2* haploinsufficiency in the absence of *CTR1* and EIN2 dosage-dependent ethylene signaling (22). This CTR1-independent regulation can be explained by our previous finding that CTR1 promotes receptor signal output, which in turn prevents EIN2 activation. In this model, the histidine kinase (HK) domain of ETR1 inhibits receptor signaling (9, 23), as evidenced by the reversion of the *ctr1-1* mutant phenotype by the *ETR1^1-349^* transgene (21).

Ethylene receptor signaling is cooperative (26, 27); we first tested whether ETR1^1-349^ receptor signaling is mediated via other receptor isoforms in the absence of CTR1. The *ers1-2*, *ers1-3*, and *ers2-3* mutations result from T-DNA insertion in the Wassilewskija ecotype, and the *etr1-10*, *etr2-3*, and *ein4-4* point mutations in the Col-0 ecotype (8, 9, 21, 24, 28). To avoid unexpected outcomes due to mixed ecotype backgrounds or T-DNA insertions at unknown loci, the mutants used in this study were all in the Col-0 ecotype background involving genetic crossings. *ERS1* and *ERS2* were each edited by the CRISPR-Cas9 technique, with the transgene crossed out by backcrossing with wild type (Col-0). The *etr1-10* mutation was previously reported, with W74* early termination (21), and the *etr2-3* and *ein4-4* mutations result in the W312* and W138* early terminations, respectively (24). Nucleotide thymine (T) insertions at positions 153 and 343 were detected in the *ers1^91*^* and *ers2^146*^* early-termination mutants, respectively (Fig. 2A). We did not isolate the *ctr1-3 etr1-10 ers1^91*^* mutant; nevertheless, compared to the extreme growth inhibition of *ctr1-3* and *etr1-10 ers1^91*^* plants, the substantially larger rosette phenotype of *ctr1-3 etr1-10 ers1^91*^* plants expressing the *etr1-1^1-349^-FLAG* or *ETR1^1-349^-FLAG* transgene indicated repression of EIN2-activated ethylene signaling independent of CTR1 (Fig. 2B). This finding aligns with the rosette phenotype of the *etr1-10 ers1^91*^* mutant, *ers1^91*^ etr2-3 ein4-4 ers2^146*^* quadruple mutant (designated *quad*), *etr1-10 ers1^91*^ etr2-3 ein4-4 ers2^146*^*quintuple mutant (designated *quint*), or *ctr1-3 etr1-10 ers1^91*^ etr2-3 ein4-4 ers2^146*^* (designated *ctr1-3 quint*) sextuple mutant that expressed the *ETR1^1-349^-FLAG* transgene (Fig. 2B). The ETR1 receptor N-terminus is sufficient to prevent EIN2-activated ethylene signaling independent of CTR1 and other receptor isoforms.

**Figure 2.**
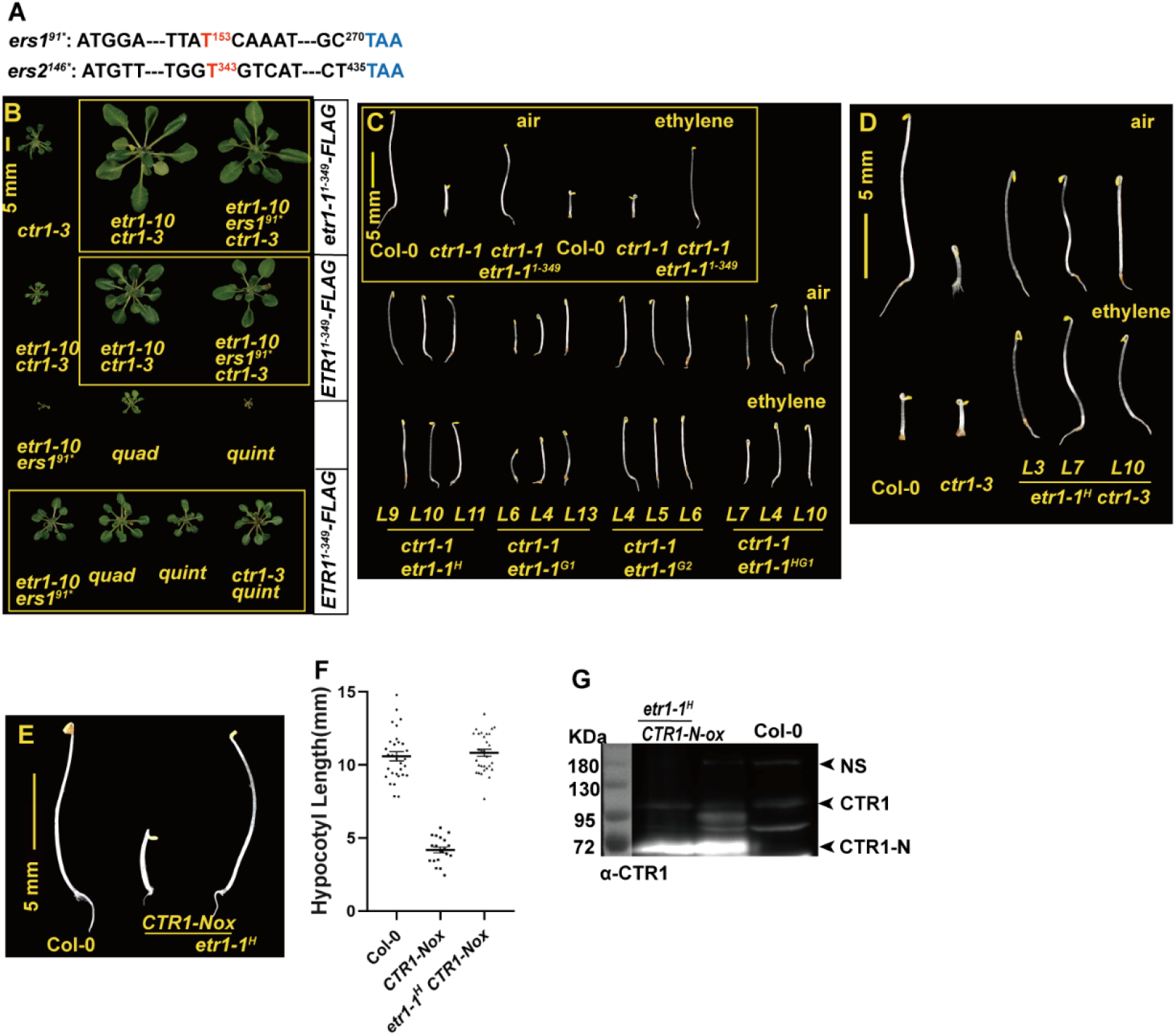
Mutations in the histidine kinase domain of ETR1 reverse the *ctr1* mutant phenotype. (A) Nucleotide sequences flanking the *ers1^91*^* and *ers2^146*^* mutants. The thymine (T) insertions at positions 153 and 343 of the *ERS1* and *ERS2* coding regions, respectively, are indicated. Dashed lines are the sequences not shown. The numbers indicate the nucleotide positions. (B) The *etr1-1^1-349^* or *ETR1^1-349^* transgene reversed the growth inhibition phenotypes of the boxed mutants carrying the *ctr1-3* allele. *Quad* and *quint* represent the *ers1^91*^ etr2-3 ein4-4 ers2^146*^* and *etr1-10 ers1^91*^ etr2-3 ein4-4 ers2^146*^* quintuple mutants, respectively. (C) The phenotype of etiolated seedlings expressing the indicated *ETR1* transgene without (air) or with ethylene treatment. (D) The *etr1-1^H^* transgene reversed *ctr1-3* seedling growth inhibition and conferred ethylene insensitivity. *CTR1-NOX*-mediated growth inhibition was prevented by the *etr1-1^H^* transgene (E), as supported by the seedling hypocotyl measurements (F). (G) Western blots showing the chemiluminescence of the CTR1 or CTR1-N immune signals of the indicated genotypes. The original image can be found from Figure2G_course_data_1 and Figure2G_course_data_2. NS: nonspecific noise. The data are presented as the means and standard errors (SEs), and the dots indicate individual measurements. The *etr1-1^H^* transgene is driven by the *ETR1* promoter. H: H353Q, G1: G515A G515A, and G2: G545A G547A.

The ETR1^1-349^ fragment does not contain the HK domain; thus, HK autophosphorylation is impaired. We tested whether mutations impairing ETR1 autophosphorylation prevent EIN2-activated ethylene signaling independent of CTR1. The H353 residue and the G1 and G2 boxes are required for ETR1 autophosphorylation. Mutations of these signature motifs [H353Q (H), G515A G517A (G1), and G545A G547A (G2)] that impair kinase activity do not impact ETR1 functionality (8, 29, 30). Similar to the effect of the *etr1-1^1-349^* transgene on *ctr1-1* seedling growth, the expression of the *etr1-1^H^*, *etr1-1^G1^*, *etr1-1^G2^*, or *etr1-1^HG1^* transgene reversed *ctr1-1* growth inhibition to various degrees (Fig. 2C). The strong *ctr1-3* mutation was also suppressed by the *etr1-1^H^* transgene (Fig. 2D). In addition, the inhibitory effect of the excess CTR1 N-terminus (CTR1-N) on ETR1 receptor signaling was fully prevented by the *etr1-1^H^* transgene (11, 22) (Fig. 2E and 2F), with a *P* value of 0.54 for the sample mean comparison between wild-type (Col-0) and *etr1-1^H^ CTR1-Nox* seedlings (Student’s *t-*test). Western blots revealed similar CTR1-N levels in *CTR1-Nox* and *etr1-1^H^ CTR1-Nox* plants (Fig. 2G).

The reversal of the *ctr1* mutant phenotype by these ETR1 variants implies that the HK domain plays an inhibitory role in ETR1 receptor signaling. Alternatively, these mutations can alter the ETR1 protein to a conformation that prevents EIN2 activation independent of CTR1 (9). The reversion of the *ctr1 ein2/EIN2* sesquimutant phenotype but not the *ctr1* mutant phenotype by *etr1-1* or *RTE1* overexpression suggested that ethylene signaling activated by a low dosage of EIN2 can be prevented by basal-level receptor signaling. In contrast, ethylene receptors are insufficient to prevent ethylene signaling when there is excess EIN2, even in the presence of CTR1 (3, 18).

### The *etp1 etp2* Double Mutation Has Little Impact on the Ethylene Response

EIN2 phosphorylation is followed by ETP1- and ETP2-mediated degradation, implying an association between EIN2 stability and phosphorylation (2–4). On the other hand, ethylene-induced EIN2 accumulation (2) does not align with the report of ethylene-induced EIN2 reduction, which is coupled with ethylene-induced EIN2 cleavage and transport to the nucleus (5). Resolving this contradiction may help us address EIN2 inactivation by ETR1 in the absence of CTR1-mediated phosphorylation. Our first task was to characterize the ethylene responsiveness on the loss of *ETP1* and *ETP2*, which was previously investigated by *amiRNA* expression but not double mutation (2).

The *ETP1* (AT3G18980) and *ETP2* (AT3G18910) loci are closely linked. We created the *etp1 etp2* double mutant using the CRISPR-Cas9 technique by sequentially editing the *ETP1* and *ETP2* genes and crossing out of the transgene (Fig. 3A). There was no distinguishable difference in seedling (Fig. 3B) or rosette (Fig. 3C) phenotypes between wild type (Col-0) and the mutants. The ethylene responsiveness of the *etp1^7*^ etp2^41*-1^* double mutant (designated *etp1 etp2*) was characterized by measuring seedling hypocotyl growth over a wide range of ethylene concentrations, and wild type (Col-0) and the double mutant exhibited identical ethylene dose-response curves (Fig. 3D).

**Figure 3.**
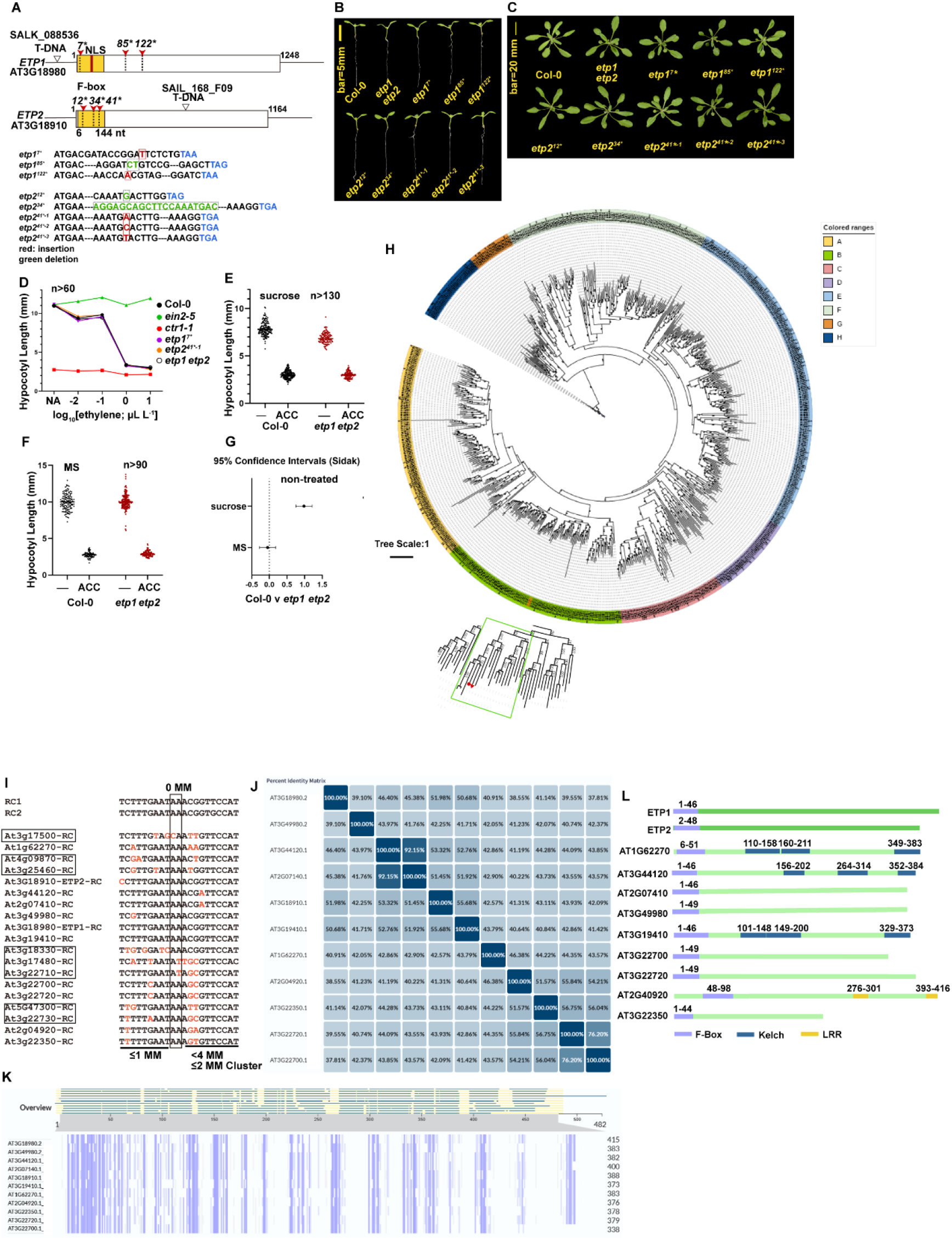
The *etp1 etp2* double mutation has little impact on the ethylene response and plant growth. (A) The gene structures of *ETP1* and *ETP2*. The F-box-coding region is boxed, where an NLS is identified for ETP1. The reported T-DNA insertion mutations are indicated by triangles. Red arrowheads denote mutations isolated from genetic editing. The dashed line-boxed nucleotide sequences indicate deletion and the solid line-boxed sequences indicate insertions. The phenotypes of the indicated *etp1 etp2*, *etp1*, and *etp2* mutants at the seedling (B) and rosette (C) stages. In this study, *etp1^7*^ etp2^41*-1^* was designated *etp1 etp2*. (D) The ethylene dose-response curves are identical for wild-type (Col-0), *etp1*, *etp2*, and *etp1 etp2* seedlings. (E) The hypocotyl measurements for the seedlings grown on the medium supplemented with sucrose. (F) Hypocotyl measurements of wild-type (Col-0) and *etp1 etp2* seedlings without sucrose supplementation. (G) Hypocotyls were longer for non-treated seedlings grown on the medium without sucrose supplementation than those with sucrose supplementation between wild type (Col-0) and the *etp1 etp2* mutant. ACC: 1-aminocyclopropane-1-carboxylic acid. (H) The phylogeny of the 745 nonredundant F-box-containing proteins, and the ETP-like subclade is boxed. The solid circles and triangles denote ETP1 and ETP2, respectively. (I) The reverse complement (RC) sequence of the amiRNA and the potentially targeted *ETP*-like genes. RC1 and RC2 are the reported amiRNA sequences (2). Mismatched sequences are shown in red. The criteria for amiRNA targeting were as follows: mismatch (MM) not allowed for nucleotides 10 and 11 as the cleavage site, up to four mismatches between positions 13 and 21 (with no more than two consecutive mismatches), and at most one mismatch allowed between positions 2 and 12 (12). The *ETP*-like genes that do not fit the criteria are boxed. The sequence identity matrix (J) and sequence alignment (K) of the 11 ETP-related proteins. The graphs were generated by UniProt (1). (L) The annotated protein structures for ETP1, ETP2, and ETP-related proteins (UniProt; https://www.uniprot.org/).

Previous studies on *ETP1* and *ETP2* involved 1-aminocyclopropane-1-carboxylate (ACC) treatment as a proxy for ethylene on a growth medium supplemented with sucrose (2). ACC is the immediate precursor of ethylene and has roles in plant growth independent of ethylene (31, 32). We investigated whether the *etp1 etp2* mutant might exhibit seedling growth inhibition in response to ACC. When supplemented with sucrose, the nontreated wild-type plants had slightly longer (by 1 mm) seedling hypocotyls than did the double mutant plants, and seedling growth was identical for both genotypes with the ACC treatment (Fig. 3E). Given that sucrose supplementation affects seedling hypocotyl growth (33), we measured seedling hypocotyl growth without sucrose supplementation, and the double mutation had little effect on growth regardless of the ACC treatment (Fig. 3F and 3G). These measurements revealed that the growth of nontreated seedling hypocotyls was inhibited by sucrose supplementation.

F-box-containing proteins constitute a superfamily, and the biological functions of a large fraction of the family members have yet to be determined. The distinct ethylene response phenotypes of the *etp1 etp2* mutant and *amiRNA* expression lines could be attributed to off-target effects of the amiRNAs. In a phylogenetic analysis of the 745 annotated F-box-containing proteins (34)(Fig. 3H and Supplemental Figure S1; with the redundant sequences removed), 19 proteins, including ETP1/ETP2, were classified into the same subclade. By sequence analysis, 11 out of the 19 *ETP1/ETP2*-related genes could be targeted by the amiRNAs (Fig. 3I). These 11 ETP-related proteins share various degrees of sequence identity, and ETP1 and ETP2 share 51.98% sequence identity (Fig. 3J and 3K). ETP1, ETP2, AT2G07410, AT3G49980, AT2G22700, AT2G22720, and AT3522350 are predicted to contain F-boxes without known structures or motifs (F-box only or FBXO). AT1G62270, AT3G44120, and AT3G19410 are predicted to be Kelch repeat F-Box proteins (KFPs), and AT2G40920 is predicted to be a leucine-rich repeat (LRR) F-Box protein (Fig. 3L).

Our present data show little impact on the ethylene response by the *etp1 etp2* double mutation, and the molecular model that EIN2 is regulated at the protein level by the two F-box proteins has yet to be reconfirmed. We were unable to reproduce the constitutive ethylene response phenotype by *amiRNA* expression (Supplemental Figure S2). It is beyond the scope of this study to determine whether possible off-target effects of amiRNAs could result in combinatorial effects that affect ethylene signaling.

### Influence of the *mhl1 mhl2* Double Mutation on Growth

EIN2 stability requires MHL1/MHL2. The *mhl1 mhl2* double mutation, which involves genetic crossing of the Col-0 and L*er* backgrounds of mutants resulting from T-DNA insertion (*mhl1*) and the *Ds* transposon (*mhl2*), reverses the *ctr1-1* mutant phenotype to the wild-type level (13). This finding implies that the stability of EIN2, which is not phosphorylated by CTR1, requires MHL1/MHL2 (13), suggesting that ETR1-mediated EIN2 inactivation in the absence of CTR1 involves MHL1/MHL2.

In contrast, we were unable to reproduce the prominent *mhl1 mhl2* ethylene-insensitive phenotype by Ma *et al.* involving the identical germplasm (13) (Supplemental Figure S3 for the seedling phenotype). Without ethylene treatment, the hypocotyls of wild-type (Col-0) seedlings were approximately 4 mm and 6 mm longer than those of *mhl1 mhl2* and *mhl1 mhl2 ctr1-1* mutant plants, respectively (Fig. 4A). With ethylene, the seedling hypocotyl of wild type was marginally shorter than that of the double or triple mutant (<0.5 mm, Fig. 4A). The previous study on *MHL1*/*MHL2* involved sucrose supplementation (13). However, even with sucrose, the *mhl1 mhl2* and *mhl1 mhl2 ctr1-1* mutants did not display the described prominent ethylene insensitivity (13). The two nontreated mutants produced shorter seedling hypocotyls than did the nontreated wild type (Col-0) by approximately 3–3.5 mm, albeit a marginally longer hypocotyl than did the wild type by approximately 0.5–0.7 mm in response to ethylene (Fig. 4B).

**Figure 4.**
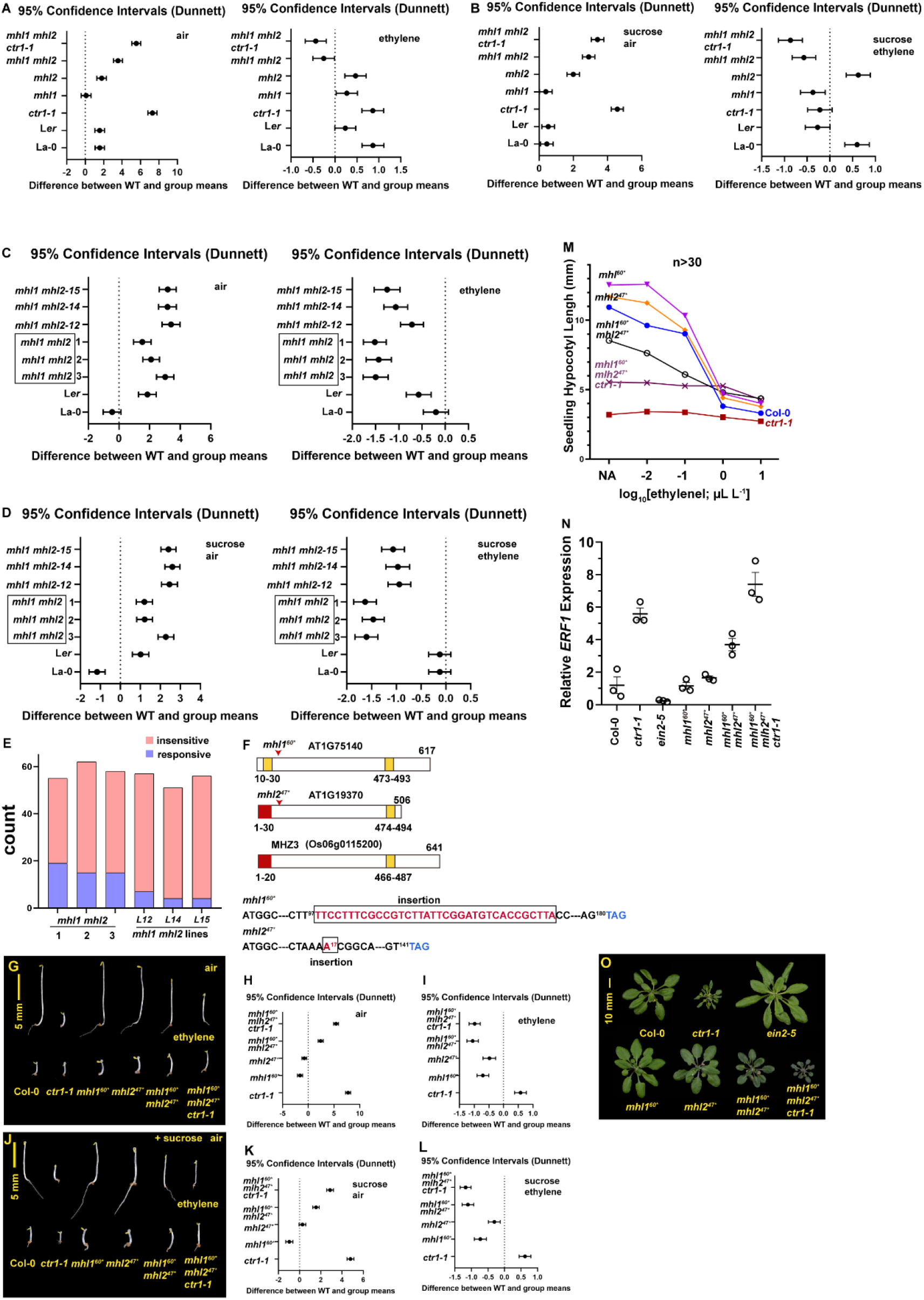
The *mhl1 mhl2* double mutation has complex impacts on growth. The mean difference (CI_0.95_) in seedling hypocotyl length between wild type (Col-0) and the indicated genotypes, without (air) or with ethylene treatment, in the absence (A) or presence (B) of sucrose supplementation. The mean difference (CI_0.95_) in seedling hypocotyl length between wild type (Col-0) and the indicated *mhl1 mhl2* double mutant lines without (air) or with ethylene treatment in the absence (C) or presence (D) of sucrose supplementation. The three boxed *mhl1 mhl2* mutant lines (denoted with the numbers 1, 2, and 3) are the measurements from three independent progenies of the reported *mhl1 mhl2* germplasm in (A) (11). (E) Number of ethylene-responsive and ethylene-insensitive roots of the indicated *mhl1 mhl2* lines. The numbers (1, 2, and 3) indicate the three independent progeny lines from the reported *mhl1 mhl2* germplasm in (A). (F) The protein structures of MHL1, MHL2, and rice MHZ3. The predicted signal peptides (red boxes) and transmembrane domains (yellow boxes) are indicated. The *mhl1^60*^* and *mhl2^47*^* mutations are denoted. Phenotypes of the *mhl1^60*^*, *mhl2^47*^*, *mhl1^60*^ mhl2^47*^*, and *mhl1^60*^ mhl2^47*^ ctr1-1* seedlings without sucrose supplementation (G) and the mean differences (CI_0.95_) in seedling hypocotyl length between the wild type (Col-0) and the indicated genotypes without (air) (H) and with (I) ethylene treatment. Phenotypes of the *mhl1^60*^*, *mhl2^47*^*, *mhl1^60*^ mhl2^47*^*, and *mhl1^60*^ mhl2^47*^ ctr1-1* seedlings subjected to sucrose supplementation (J) and the mean differences (CI_0.95_) in seedling hypocotyl length between the wild type (Col-0) and the indicated genotypes without (air) (K) and with (L) ethylene treatment. The sample size was greater than 30 for each measurement. Ethylene dose-response curves (*n*≥30) (M) and RT-qPCR measurements of the *ERF1* level (N) in wild-type (Col-0), *mhl1^60*^*, *mhl2^47*^*, *mhl1^60*^ mhl2^47*^*, and *mhl1^60*^ mhl2^47*^ ctr1-1* plants. Circles are independent RT-qPCR measures, and the data are presented as the means and SEs. (O) The rosette phenotype of the wild type and the indicated mutants.

In addition to the reported *mhl1 mhl2* germline, analysis of independent *mhl1 mhl2* double mutant lines from the cross revealed 2-3 mm growth inhibition, compared to wild type, in the absence of ethylene (Fig. 4C). These mutants exhibited growth inhibition and did not display the reported extent of ethylene insensitivity, with sucrose supplementation (Fig. 4D). This prompted us to re-examine the reported *ein2*-level ethylene-insensitive root phenotype (13). Ethylene-treated etiolated seedlings of these *mhl1 mhl2* lines produced both ethylene-insensitive and -sensitive root phenotypes on sucrose-supplemented media (Fig. 4E and Supplemental Figure S4). Notably, the root phenotype only appeared on media supplemented with sucrose (Supplemental Figure S3). The reported ethylene-insensitive root phenotype is likely associated with other loci and is sucrose-dependent.

We considered the possibility that the reported *mhl1 mhl2* double mutant generated by genetic crossing involving two ecotypes and T-DNA/*Ds* mutations (13) could produce complex phenotypes independent of the ethylene phenotype. To minimize genetic background noise, we generated the *mhl1^60*^*, *mhl2^47*^*, and *mhl1^60*^ mhl2^47*^* mutants using the CRISPR-Cas9 technique involving the Col-0 ecotype, and the transgene was crossed out (Fig. 4F). Without ethylene treatment, wild-type (Col-0) plants produced much longer seedling hypocotyls than did the *mhl1^60*^ mhl2^47*^* and *mhl1^60*^ mhl2^47*^ ctr1-1* mutants by approximately 2 mm and 5 mm, respectively (Fig. 4G and 4H). With ethylene, the seedling hypocotyls of the two mutants were slightly longer than those of the wild type (Col-0) by approximately 1.2 mm (Fig. 4G and 4I). In contrast to the sucrose-dependent ethylene-insensitive root phenotype of the T-DNA/*Ds*-insertional *mhl1 mhl2* mutant (Supplemental Figure S4), the *mhl1^60*^ mhl2^47*^* mutation did not confer ethylene insensitivity to the root in the presence of sucrose supplementation (Fig. 4J). Sucrose supplementation reduced seedling hypocotyl length, with hypocotyl length differences of approximately 1.8 mm and 3 mm between the wild-type (Col-0) and *mhl1^60*^ mhl2^47*^* and *mhl1^60*^ mhl2^47*^ ctr1-1* seedlings, respectively (Fig. 4J and 4K). Similarly, after ethylene treatment, the hypocotyls of the *mhl1^60*^ mhl2^47*^* or *mhl1^60*^ mhl2^47*^ ctr1-1* seedlings were longer than those of the wild type (Col-0) plants by approximately >1 mm (Fig. 4J and 4L).

An ethylene dose-response assay revealed markedly shorter hypocotyls for *mhl1^60*^ mhl2^47*^ ctr1-1* seedlings than for wild-type seedlings at low ethylene concentrations (0, 0.01, and 0.1 µL L^-1^) (Fig. 4M). This pronounced growth inhibition at low ethylene concentrations cannot be simply attributed to the loss of EIN2 stability, which is consistent with the RT‒qPCR results revealing elevated levels of ethylene-induced *ETHYLENE RESPONSE FACTOR1* (*ERF1*) in the *mhl1^60*^ mhl2^47*^* and *mhl1^60*^ mhl2^47*^ ctr1-1* mutants independent of ethylene treatment (Fig. 4N). In addition to the complex ethylene-independent seedling growth inhibition phenotype, the *mhl1^60*^ mhl2^47*^* double mutation did not reverse the *ctr1-1* mutant phenotype, and the double or triple mutant also exhibited growth inhibition at the rosette stage (Fig. 4O).

Although the degree of ethylene-induced seedling hypocotyl growth inhibition was marginally weaker for the *mhl1 mhl2* mutant than for wild type, *ERF1* induction was not impacted, and the mutant roots did not exhibit ethylene insensitivity. Our study revealed that the reported ethylene-insensitive root phenotype was sucrose-dependent and independent of the *mhl1 mhl2* mutation, indicative of other loci responsible for the root phenotype. Our present data did not support that ethylene signaling is impacted by the *mhl1 mhl2* double mutation; rather, the double mutation may impact growth throughout development.

### Subcellular Localization of ETP1/ETP2 and MHL1/MHL2 and Their Effects on Ethylene Signaling

Our genetic studies do not support the involvement of ETP1/ETP2 in ethylene signaling. Nevertheless, *ETP1/ETP2* overexpression changes the ACC response phenotype to various degrees (2). These findings prompted us to investigate the subcellular localization of ETP1/ETP2 in Arabidopsis and their effects on ethylene signaling.

Yeast two-hybrid assays determine the interaction of ETP1/ETP2 with the carboxyl portion of EIN2, and the exact subcellular compartment where the interaction occurs has yet to be determined (2). In cells of transgenic Arabidopsis plants, eYFP-tagged ETP1 was localized to the nucleus, and ETP2 was localized to the cytoplasm (Fig. 5A and 5B), which is in line with the annotated nuclear localization signal (NLS) for ETP1 but not for ETP2 (Fig. 3A). Unlike nuclear EIN2 speckle formation, ETP1 was prevalent in the nucleus. The colocalization study for EIN2-mRFP and ETP1-eYFP or ETP2-eYFP was unsuccessful in transgenic Arabidopsis plants, likely due to transgene silencing. With transient expression in Arabidopsis protoplasts, EIN2-mRFP and ETP1-eYFP displayed distinct fluorescence patterns and distributions in the nucleus, in line with their distinct fluorescence patterns in the cells of transgenic Arabidopsis plants. Whether the two proteins may interact in part or colocalize needs further investigation. EIN2-mRFP and ETP2-eYFP separated from each other in the cytoplasm of Arabidopsis protoplasts, indicating the absence of interactions *in vivo* (Fig. 5C and 5D).

**Figure 5.**
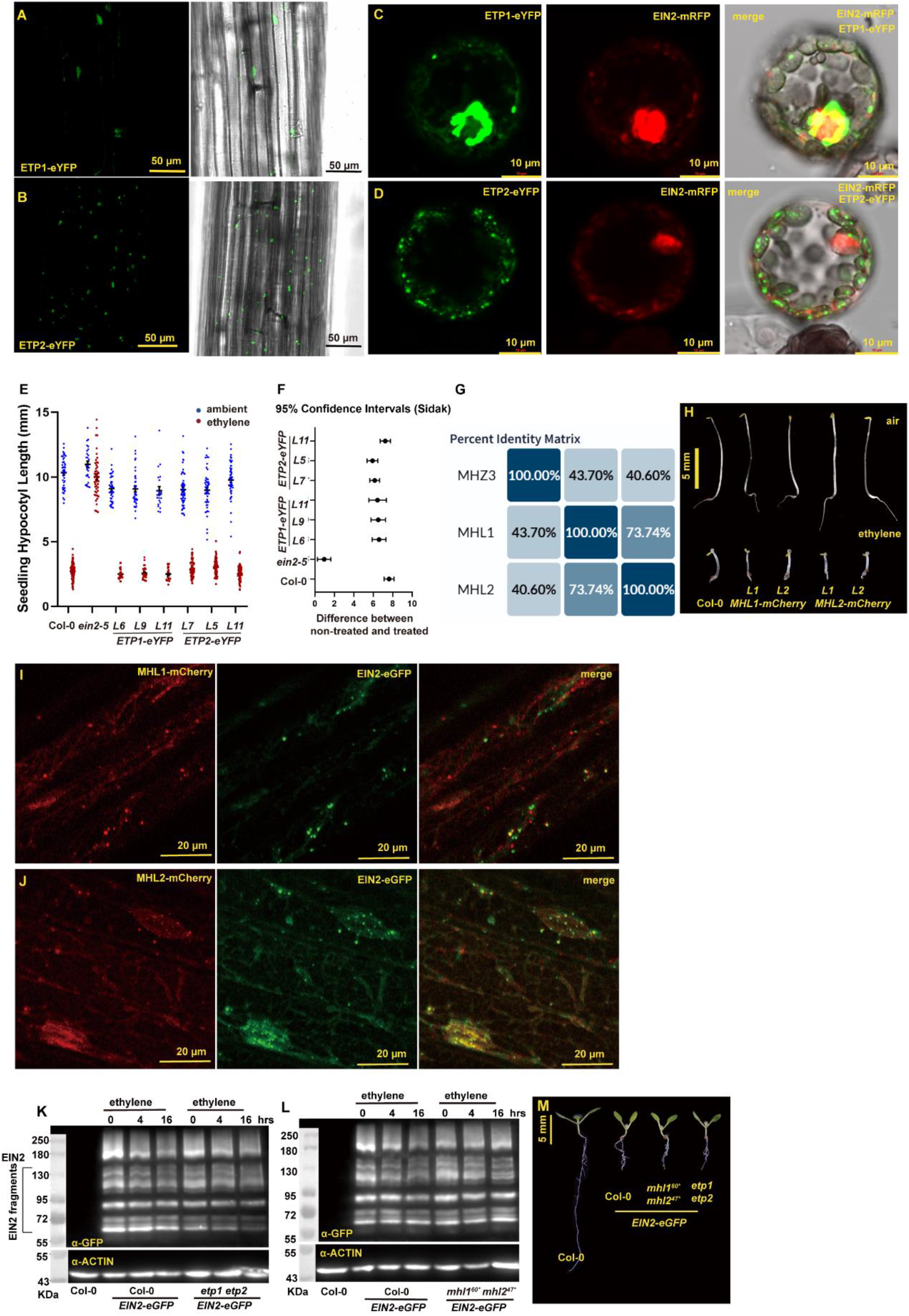
The subcellular localization of ETP1/ETP2 and MHL1/MHL2 and their effect on EIN2. Fluorescence of ETP1-eYFP (A) and ETP2-eYFP (B) in cells of transgenic Arabidopsis plants involving the *UBQ10* promoter. The fluorescence for EIN2-mRFP and eYFP-fused ETP1 (C) and ETP2 (D) in Arabidopsis protoplasts. The *35S* promoter was involved in transient expression in Arabidopsis protoplasts. Seeding hypocotyl measurements of the indicated *ETP1-eYFP* or *ETP2-eYFP* overexpression lines (E) and the mean differences (CI_0.95_) between the nontreated and ethylene-treated seedlings (F). The sample size was greater than 30 seedlings for each measurement, and the dots indicate individual seedling measurements. The data are presented as the means and standard errors (SEs). (G) The sequence identity matrix of MHL1, MHL2, and rice OsMHZ3, and the graph was generated by UniProt (42). (H) The phenotype of etiolated seedlings of the indicated *MHL1* or *MHL2* overexpression lines without (air) and with ethylene treatment. The fluorescence of *MHL1-mCherry* (I) and *MHL2-mCherry* (J) coexpressing *EIN2-eGFP* in cells of transgenic Arabidopsis plants. (K) and (L) Western blots showing the chemiluminescence of EIN2-eGFP with the indicated genotypes expressing the *EIN2-eGFP* transgene, which was genetically introduced into the double mutants. The original images can be found from Figure5K/L_source_data_1 and Figure5K/L_source_data_2. The *UBQ10* promoter was used for the *MHL1/MHL2/ETP1/ETP2* transgenes, and the *SUPER* promoter was used for the *EIN2-eGFP* transgene. (M) The phenotype of wild-type, *mhl1 mhl2*, or *etp1 etp2* seedlings expressing the *EIN2-eGFP* transgene.

The distinct localizations of ETP1 and ETP2 imply that their molecular functions differ. Qiao *et al.* reported that the hypocotyl growth of *35S:ETP1* seedlings was inhibited to a lesser degree by ACC than that of wild-type plants (Col-0), and root growth was completely inhibited. Although the measurements appeared to be incorrect (Fig. 5D by Qiao *et al.*), ACC-treated *ETP2*-overexpressing plants produced slightly longer seedling hypocotyls and roots than did ACC-treated wild-type plants (Col-0) (2). The distinct effects of *ETP1* and *ETP2* overexpression on seedling hypocotyl and root growth also imply that the two proteins have different functions. We examined the ethylene response phenotypes of our *ETP1-eYFP* and *ETP2-eYFP* transgenic plants, which were fully ethylene responsive (Fig. 5E and 5F).

Ethylene promotes coleoptile growth and inhibits root elongation of rice seedlings (13, 35, 36). Rice *OsMHZ3* overexpression results in a moderate degree of root growth inhibition, by approximately 30%, with coleoptile elongation being unaffected (13). The possibility of excess MHL1/MHL2 leading to ethylene activation was tested. MHL1 and MHL2 share 73.74% protein sequence identity; MHL1 is predicted to have two transmembrane domains, and MHL2 has a signal peptide and a transmembrane domain (Figs. 4F and 5G), which agrees with the AlphaFold prediction that MHL2 but not MHL1 is structurally similar to rice OsMHZ3 (https://alphafold.ebi.ac.uk/entry/Q9FRK5)(37, 38). As expected, *MHL2* but not *MHL1* overexpression had an effect on ethylene signaling similar to that of *OsMHZ3* overexpression. We first constructed transgenic Arabidopsis plants expressing *MHL1-mCherry* or *MHL2-mCherry*. Etiolated seedlings expressing the transgenes were phenotypically similar to wild-type (Col-0) plants in the absence or presence of ethylene (Fig. 5H). Apart from their partial colocalization at the ER-like network, transgenic Arabidopsis plants coexpressing *EIN2-eGFP* with either *MHL1-mCherry* or *MHL2-mCherry* exhibited separation of EIN2 particles from MHL1, and EIN2 particles appeared to overlap with MHL2 particles (Fig. 5I and 5J). Although the present results indicate that EIN2 and MHL2 are colocalized, this colocalization is insufficient to explain the biological significance of ethylene signaling.

Our genetic and transgenic studies showed little effect of the loss or overexpression of *ETP1/ETP2* and *MHL2/MHL2* on ethylene signaling, suggesting that EIN2 levels are regulated independently of the two protein sets. This was next tested involving an *EIN2* overexpressor that exhibited a moderate degree of constitutive ethylene signaling. In contrast to the finding that amiRNA that downregulates *ETP1/ETP2* levels substantially increases EIN2 levels (2), western blot analysis revealed that the level and cleavage of EIN2-eYFP were unaffected regardless of ethylene treatment in the *etp1 etp2* mutant (Fig. 5K). Similarly, the *mhl1 mhl2* mutation had little effect on the EIN2-eYFP level and cleavage (Fig. 5L). This is corroborated by the similar growth phenotypes of *EIN2-eYFP*, *EIN2-eYFP etp1 etp2*, and *EIN2-eYFP mhl1 mhl2* seedlings (Fig. 5M), where the *EIN2-eYFP* transgene was introduced by genetic crossing and thus expressed at the same transgene locus.

### Ethylene Receptors and EIN2 Associate with the Rough Endoplasmic Reticulum

Our present data supported that ETP1/ETP2 or MHL1/MHL2 are minimally involved in ethylene signaling, suggesting that the repression of EIN2-activated ethylene signaling by the ETR1 receptor independent of CTR1 may occur at other levels.

We first investigated the subcellular localization of EIN2. Western blots detected EIN2 and the fragments primarily in the membrane fraction, a trace amount in the nuclear fraction, and not detected in the cytoplasmic soluble fraction. The localization and complexity of EIN2 were unaffected by ethylene treatment (Fig. 6A). A small amount of membrane contamination, marked by the ER lumen protein BiP, was detected in the nuclear fraction; we cannot exclude the possibility that the trace amount of nuclear EIN2 was from the membrane. The membrane localization of EIN2 and EIN2 fragments aligned with those of independent reports (13, 18, 19). The membrane fraction was subjected to sucrose density gradient fractionation. Western blots detected EIN2, EIN2 fragments, and CTR1 from the same fractions, with a density shift upon Mg^2+^ level elevation, which marks the rough ER fractions (Fig. 6B) (39, 40). This finding also aligns with the report that CTR1 associates with the rough ER (39).

**Figure 6.**
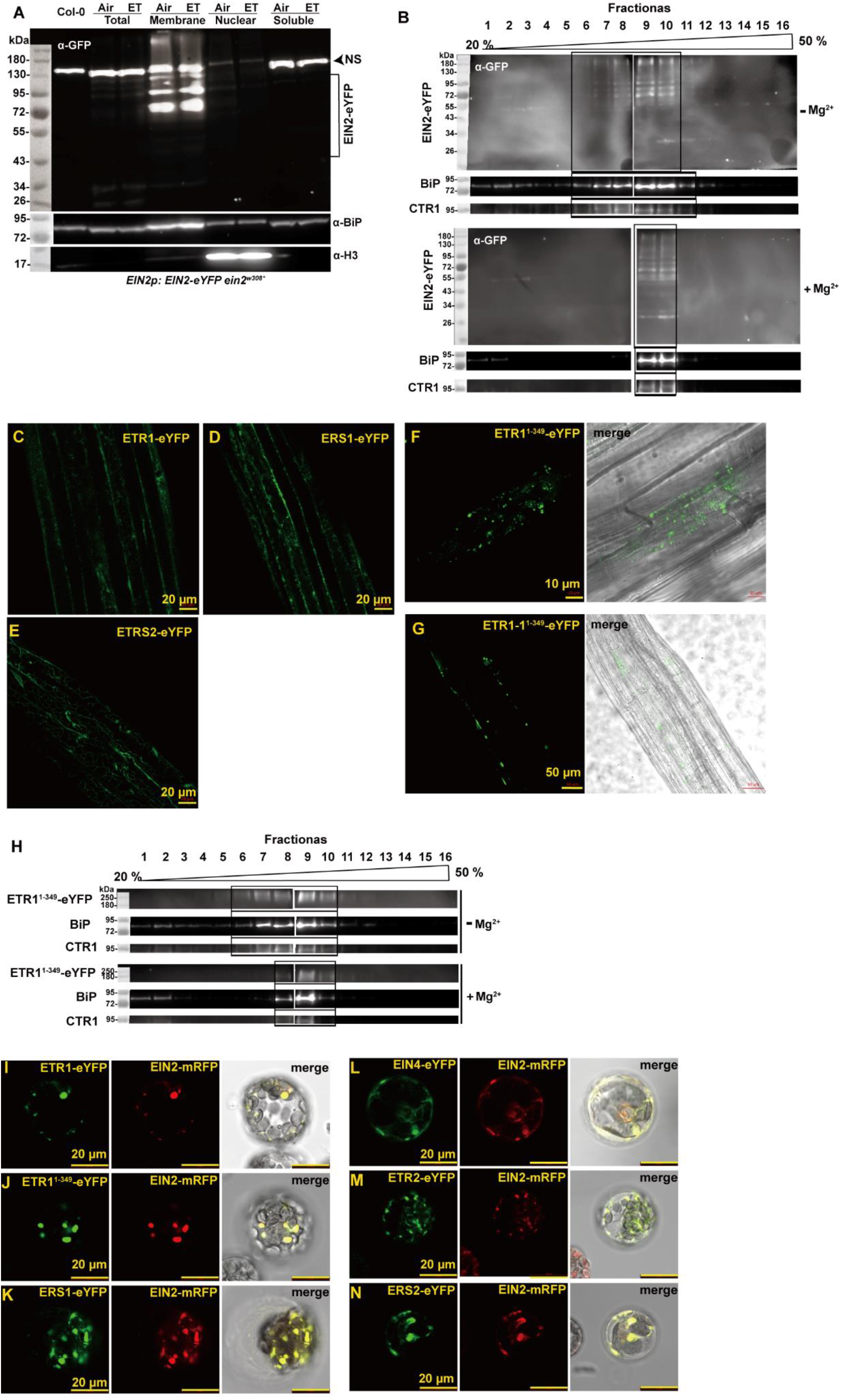
Subcellular localization of EIN2, ETR1, and CTR1. (A) Western blots showing the chemiluminescence of the eYFP-fused EIN2 expressed in the *ein2^W308*^* mutant. Total: total protein, Membrane: membrane fraction, Nuclear: nuclear fraction, and Soluble: cytoplasmic soluble fraction. NS: nonspecific immune signals. BiP is an endoplasmic reticulum (ER) protein used as a marker for membrane proteins. H3 is the nuclear HISTONE3 protein as a marker for nuclear proteins. NS: nonspecific immune signals. (B) Western blots revealing the chemiluminescence of the eYFP-fused EIN2 protein from the membrane fractionated by a sucrose density gradient (20% - 50% sucrose) under conditions of low (- Mg^2+^) and elevated (+Mg^2+^) magnesium levels. The same fractions were also probed with CTR1 antibodies to determine the ER-associated CTR1 protein. The fractions enriched with EIN2-eYFP or CTR1 are boxed. Fractions 1-8 and 9-16 were separated on independent gels, and the two western blots were separated by space. (C)-(E) Fluorescence of the eYFP-fused ETR1, ERS1, or ERS2 proteins in cells of transgenic Arabidopsis plants. Fluorescence of the eYFP-fused ETR1^1-349^ (F) and ETR1-1^1-349^ (G) proteins in cells of transgenic Arabidopsis plants. These receptor transgenes were driven by the *UBQ10* promoter. (H) Western blots showing the chemiluminescence of the eYFP-fused ETR1^1-349^ protein from the membrane fractionated by a sucrose density gradient (20% - 50% sucrose) under conditions of low (- Mg^2+^) and elevated (+Mg^2+^) magnesium levels. The same fractions were also probed with CTR1 antibodies to determine the ER-associated CTR1 protein. Fractions 1-8 and 9-16 were separated on independent gels, and the two western blots are shown separately. Fluorescence of Arabidopsis protoplasts co-expressing EIN2-mRFP with ETR1-eYFP (I), ETR1^1-349^-eYFP (J), ERS1-eYFP (K), EIN4-eYFP (L), ETR2-eYFP (M), or ERS2-eYFP (N). The transient expression of ethylene receptor transgenes and *EIN2-mRFP* in Arabidopsis protoplasts involved the *35S* promoter. The original images for western blots can be found from Figure6A/B/H _source_data_1 and Figure6A/B/H_source_data_2.

We next investigated the subcellular localization of the ETR1 receptor. ETR1-eYFP, ERS1-eYFP, and ERS2-eYFP produced a network structure with a few speckles in cells of the transgenic Arabidopsis plants (Fig. 6C-6E). The *ETR2-eYFP* or *EIN4-eYFP* transgenic lines that produce fluorescence were not available. In contrast to the network-like structure for the ETR1-eYFP fluorescence pattern, the eYFP fluorescence of ETR1^1-349^-eYFP or ETR1-1^1-349^-eYFP in the cells of transgenic Arabidopsis was granular (Fig. 6F and 6G), similar to that of the reported EIN2 particles (18). Sucrose density gradient fractionation revealed a Mg^2+^-dependent density shift of ETR1^1-349^-eYFP and CTR1 (Fig. 6H), suggesting the association of ETR1^1-349^-eYFP with the rER. The co-localization of the ethylene receptor members and EIN2 was also supported by the transient expression of the receptor-eYFP fusion protein and EIN2-mRFP (Fig. 6I-6N). Notably, the protein fluorescence appeared as granules in Arabidopsis protoplasts, in contrast to the network structure when expressed in cells of transgenic Arabidopsis plants. The granular fluorescence patterns align with those of a previous report (18), likely resulting from ER body formation or other ER structures (41).

### The ETR1 Ethylene Receptor Interacts with EIN2

CTR1 was determined to interact with and phosphorylate EIN2 (3); however, this interaction was not detectable for transiently expressed EIN2-C and CTR1 in the tobacco epidermis (42). Given that EIN2 and the EIN2 fragments, ETR1, and CTR1 are detectable in the membrane and enriched in the rough ER fractions, we tested whether these three proteins may physically interact as a complex.

CTR1 did not coimmunoprecipitate with EIN2-eYFP in transgenic *ein2^W308*^* plants expressing *EIN2-eYFP* (Fig. 7A). In wild-type Arabidopsis protoplasts coexpressing *eYFP* or *EIN2-eYFP* with *ETR1^1-349^-3×FLAG*, *ETR1-3×FLAG*, or *ETR1^H^-3×FLAG*, the three ETR1 variants coimmunoprecipitated with EIN2-eYFP but not with eYFP (Fig. 7B and 7C). Notably, when expressed in Arabidopsis protoplasts, EIN2 was predominantly noncleaved, and it was primarily the noncleaved EIN2 that coimmunoprecipitated with ETR1 (Fig. 7B). This does not exclude the possibility that the cleaved EIN2 fragments were also associated with ETR1.

**Figure 7.**
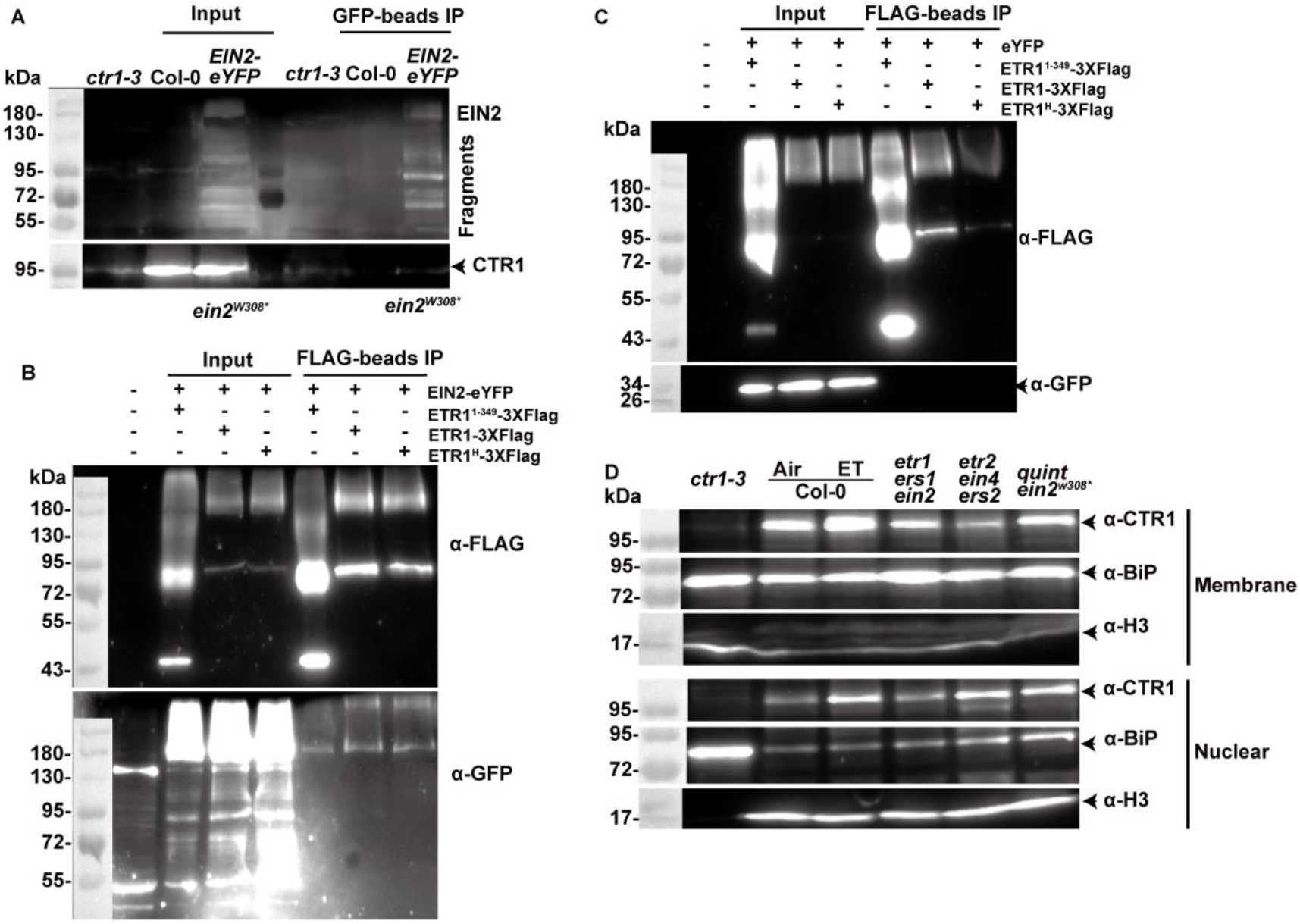
Coimmunoprecipitation studies of EIN2 with ETR1 or CTR1. (A) CTR1 was not detected from the proteins coimmunoprecipitated with EIN2-eYFP. Proteins are from the *EIN2p:EIN2-eYFP ein2^W308*^* transgenic plants. A faint nonspecific immune signal is detected from proteins of the *ctr1-3* plant (input) and the Co-IP proteins (Col-0 and *EIN2-eYFP*). (B) EIN2-eYFP can be detected from proteins coimmunoprecipitated with 3×FLAG-tagged ETR1 or the indicated variants from proteins transiently expressed in Arabidopsis protoplasts. (C) As a control, eYFP was not detected in the proteins coimmunoprecipitated with 3×FLAG-tagged ETR1 or the idicated variants from proteins transiently expressed in Arabidopsis protoplasts. (D) Western blots showing the chemiluminescence of the CTR1 protein in the membrane and nuclear fractions. Air: nontreated; ethylene: ethylene-treated. *quint ein2^W308*^*: the *etr1-10 ers1^91*^ etr2-3 ein4-3 ers2^146*^ ein2^W308*^* sextuple mutant. BiP is an endoplasmic reticulum (ER) protein used as a marker for membrane proteins. H3 is the nuclear HISTONE3 protein as a marker for nuclear proteins. The *ETR1* transgenes were driven by the *UBQ10* promoter, and *eYFP* and *EIN2-eYFP* were driven by the *35S* promoter. The original images for western blots can be found from Figure7A/B/C/D _source_data_1 and Figure7A/B/C/D_source_data_2.

CTR1 does not have a transmembrane helix, and its association with the ER is considered a result of its interaction with ethylene receptor isoforms (10, 43), providing further support for our hypothesis that CTR1 docking at ETR1 promotes ETR1 receptor signaling, which in turn inactivates EIN2. On the other hand, the receptor-dependent ER association has never been tested in receptor-free mutants. The subcellular localization of CTR1 was examined in the *etr1-10 ers1^91*^ ein2^W308*^*, *etr2-3 ein4-4 ers2^146*^*, and *etr1-10 ers1^91*^ etr2-3 ein4-4 ers2^146*^ ein2^W308*^* mutants, with wild-type (Col-0) and *ctr1-3* plants serving as controls. This helps us to investigate whether the subcellular localization of CTR1 is determined by ethylene receptors or subfamily members. Subfamily I-defective *etr1-10 ers1^91*^*and receptor-defective quintuple mutants are lethal, and mutant growth is strongly inhibited; the *ein2^W308*^* mutation rescued growth and facilitated this study. In wild type (Col-0), CTR1 was associated with the membrane and nuclear fractions, and ethylene treatment elevated the CTR1 level in both the membrane and nuclear fractions. The relative level of membrane-associated CTR1 in the *etr1-10 ers1^91*^ ein2^W308*^* mutant was greater than that in the subfamily II-defective mutant, and the nuclear CTR1 level was greater in the subfamily II-defective mutant than in the *etr1-10 ers1^91*^ ein2^W308*^* mutant. In the *etr1-10 ers1^91*^ etr2-3 ein4-4 ers2^146*^ ein2^W308*^* sextuple mutant, CTR1 was localized to the membrane and nuclear fractions. The ER that surrounds and attaches to the nucleus can be enriched in the nuclear fraction, and there appears to be a small fraction of nuclear contamination in the membrane fraction and membrane contamination in the nuclear fraction. Nevertheless, according to the level of CTR1 and the degree of contamination, the contamination did not affect the conclusion that CTR1 is associated with the two fractions (Fig. 7D). The elevated nuclear CTR1 levels in ethylene-treated wild type align with reports of ethylene-induced CTR1 transport to the nucleus (42). The association of CTR1 with the membrane fraction in those mutants suggested that the loss of subfamily II members reduces the level of membrane-bound CTR1, coupled with elevated levels of nuclear CTR1. The reduced membrane CTR1 level in the subfamily II mutant agrees with the report by Gao *et al*. (43). The association of CTR1 with the membrane fraction in the sextuple mutant suggested that CTR1 membrane localization can be independent of its interaction with receptor isoforms or EIN2.

## Discussion

The phosphorylation status, stability, cleavage, and protein transport of EIN2 are considered to be intertwined processes that dictate ethylene signaling (2, 3, 5, 13). In contrast to the present model, our study suggested that ethylene signaling activation by non-phosphorylated or phosphomimic EIN2 variants is ethylene dependent and that ETP1/ETP2 or MHL1/MHL2 has little role in ethylene signaling. The EIN2 protein, either cleaved or not cleaved, is associated with the rough ER membrane. The ethylene receptor ETR1 can repress EIN2-activated ethylene signaling independent of CTR1, and ETR1 and EIN2 physically interact. CTR1 is associated with the rough ER and the nucleus, independent of ethylene receptor isoforms or EIN2. These findings suggest other explanations for the molecular events involved in ethylene signaling activation.

Our investigation of the possible regulation of EIN2 stability involving EIN2 phosphorylation status in association with ETP1/ETP2 or MHL1/MHL2 (3–5, 13) led to different findings. The *etp1 etp2* double mutation affects ethylene signaling little, and the *mhl1 mhl2* mutation impacts growth. The distinct subcellular localizations of ETP1 and ETP2 imply that their molecular functions differ, which aligns with the different ACC-response phenotypes caused by *ETP1* and *ETP2* overexpression reported by Qiao *et al.*(2), where *ETP1* overexpression does not confer ACC insensitivity in roots. In contrast, our study revealed that *ETP1* or *ETP2* overexpression had little impact on the ethylene response phenotype. Sequence analysis suggested that the amiRNAs potentially target 11 *ETP*-like genes of the 19 ETP-like subclade members and could have complex effects on seedling growth or ACC/ethylene responses. We cannot exclude the possibility that other F-Box-encoding genes, in addition to those of the ETP-like subclade, are also targeted by amiRNAs. Unfortunately, we failed to reproduce the reported effect of *amiRNA* expression to further test this possibility. Notably, according to yeast two-hybrid assays, ETP1/ETP2 does not interact with the known 19 Arabidopsis SKP-LIKE (ASK) proteins, raising the question of whether ETP1/ETP2 is capable of SCF^ETP1/ETP2^ ubiquitin ligase complex formation to mediate EIN2 ubiquitination (44). This finding aligns with the little impact of the *etp1 etp2* double mutation on the EIN2-eGFP level and the *EIN2-eGFP* plant phenotype. In addition to its role in 26S proteasome-mediated protein degradation, protein ubiquitination as a code determines various biological activities, such as the regulation of protein interactions, activity, or localization (45). At present, it is beyond the scope of our study to extend investigations testing the role of ETP-related proteins in EIN2 ubiquitination and degradation.

Our intensive investigations revealed marginally weaker ethylene hypocotyl growth inhibition in the *mhl1 mhl2* mutant than in wild type, and mutant growth was greatly inhibited in the absence of ethylene treatment throughout growth. Seedling growth can be affected by various factors; with a marginal growth change, whether the *mhl1 mhl2* double mutation truly confers ethylene insensitivity requires further study. The shortened roots in response to ethylene, elevated *ERF1* levels, and rosette growth defect phenotype of nontreated *mhl1 mhl2 ctr1-1* plants do not support the double mutation resulting in EIN2 instability and ethylene insensitivity. The prominent ethylene insensitivity of *mhl1 mhl2* seedlings reported by Ma *et al.*(13) may be attributed to unknown loci associated with other T-DNA/*Ds* insertions or to ecotype background mixing (13). This finding aligns with the segregation of the ethylene-insensitive root phenotype, which is dependent on sucrose supplementation and not associated with the *mhl1 mhl2* mutation. Although EIN2 and MHL1/MHL2 have been experimentally determined to localize to the ER (5, 46, 47), their colocalization cannot indicate the presence of protein interactions, and lines of evidence from our study did not support their functional association. The overexpression of *MHL1* or *MHL2* had little effect on the ethylene response phenotype, providing further support for little dependence of EIN2 stability regulation on these two proteins. This finding is consistent with the unchanged EIN2-eGFP level in the *mhl1 mhl2* mutant and the finding that rice *OsMHZ3* overexpression results in moderate coleoptile growth inhibition, with root growth being unaffected (13). The possibility that *OsMHZ3* overexpression does not truly change ethylene responsiveness should be considered.

In contrast to the proposed ethylene-dependent single cleavage and nuclear transport of EIN2 (5), lines of independent evidence support complex EIN2 cleavages and subcellular localization (13, 17–20), independent of ethylene treatment or the phosphorylation residues. EIN2 or cleaved fragments are not detected in the cytoplasmic soluble fraction (19), which is consistent with its separation from membraneless processing bodies in Arabidopsis protoplasts or cells of transgenic Arabidopsis plants (18, 48). Mutations or deletions of the four phosphorylation residues do not lead to autonomous ethylene signaling, and these EIN2 variants are responsive to ethylene. These data support the argument that nonphosphorylated EIN2 is not autonomously active. In addition to the evidence from transgenic studies, the *ctr1 ein2/EIN2* sesquimutant phenotype was substantially reversed by the ethylene-insensitive *etr1-1* mutation or *RTE1* overexpression, supporting the argument that the ETR1 receptor prevents EIN2-activated ethylene signaling independent of CTR1-mediated phosphorylation.

The present molecular pathway is proposed based on the genetic framework, where CTR1 acts downstream of ethylene receptors and upstream of EIN2. Genetic analysis that determines the interaction between two genes does not define the upstream or downstream relationship of the two proteins. The *ctr1* mutation that masks the ethylene-insensitive ethylene receptor mutations may imply at least three possibilities: CTR1 acts downstream of the receptors, coordinates with the receptors, or activates the receptors. In this study, we propose that CTR1 functions as an activator of the ethylene receptor; its docking at the HK domain alleviates the inhibitory effect of the HK domain on receptor signaling (9). The deletion or mutation of the H353, G515 G517, or G545 G457 residues of the HK domain results in a conformational change that allows the receptor signal output to prevent EIN2-activated ethylene signaling in the absence of CTR1. These mutations impair ETR1 autophosphorylation (29, 30); ETR1 receptor signaling is independent of its kinase activity (8), and whether this kinase activity may affect the conformational changes of the ETR1 protein and receptor signaling has to be determined. In response to ethylene, receptor signal output is prevented, EIN2 is inhibited, and ethylene signaling is activated (Fig. 8). This model explains the weaker phenotype for the *ctr1-3* mutant than for the receptor quintuple mutant, where basal-level receptor signaling still in part prevents EIN2-activated ethylene signaling in the *ctr1-3* mutant. In the receptor quintuple mutant, which lacks basal-level receptor signaling, ethylene signaling is fully activated by EIN2. This argument is consistent with the complete reversion of the quintuple mutant phenotype by the *ein2* mutation. Notably, *RTE1* overexpression or the *etr1-1* mutation had distinct effects on the sesquimutant and *ctr1* mutant phenotypes. The distinct phenotypes reflect basal-level receptor activity on ethylene signaling, which is determined by the EIN2 dosage of the two mutants. The effect of EIN2 dosage on ethylene signaling aligns with constitutive ethylene signaling activation when the wild-type EIN2 is in excess (3, 18). Both CTR1 and EIN2 can interact with ETR1, the evidence for the CTR1-EIN2 interaction is controversial, and the association of CTR1 with the rough ER can be independent of the ethylene receptor and EIN2. There is likely a scaffold that supports proximity for the dynamic interaction or complex formation of CTR1, the ethylene receptor, and EIN2.

**Figure 8.**
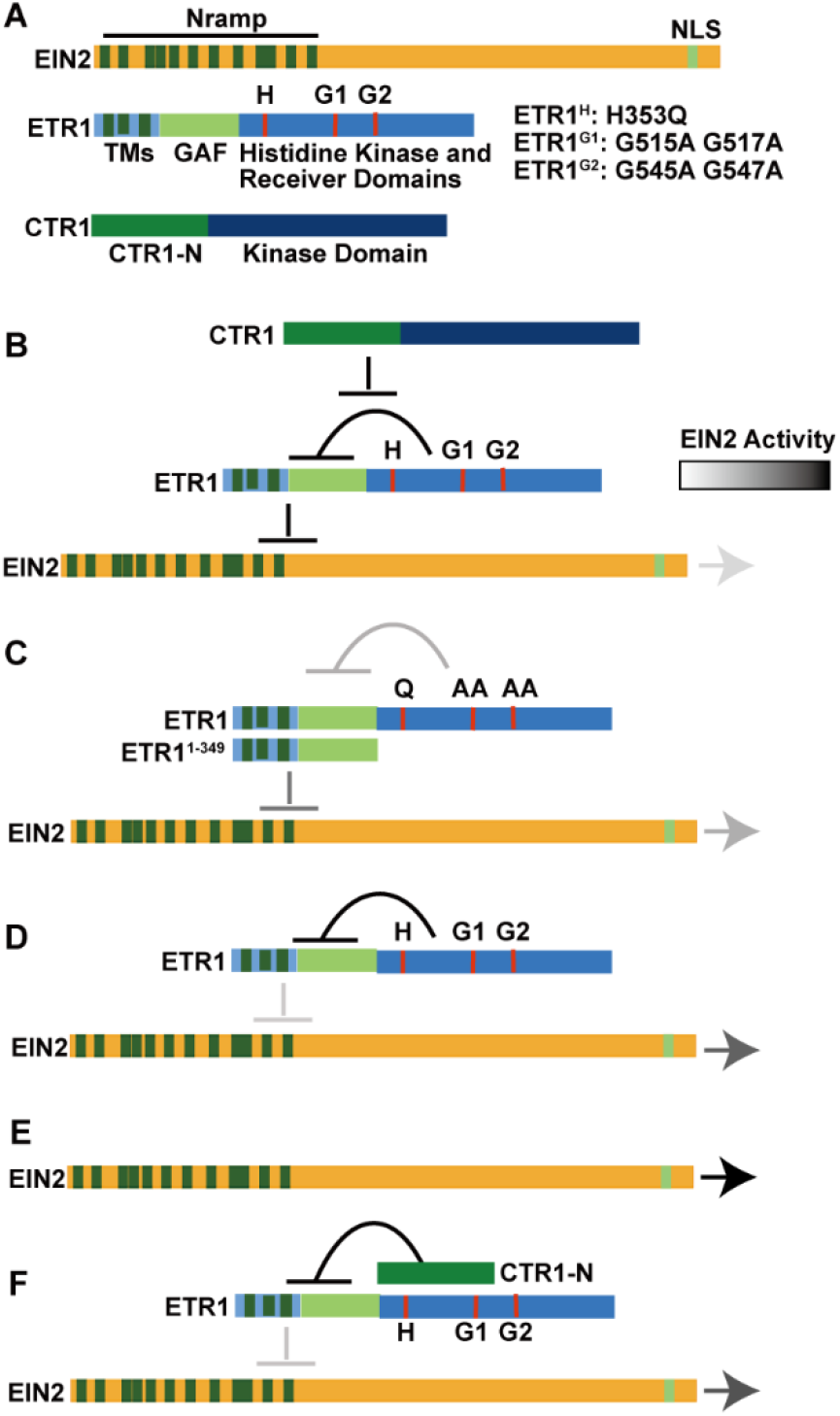
An alternative model of ethylene signaling. (A) The structures of the EIN2, ETR1, and CTR1 proteins are shown. The 12 transmembrane helices of the Nramp domain and the nuclear localization signal (NLS) of EIN2 were annotated at UniProt (1). The H353 (H), G515 G517 (G1), and G545 G547 (G2) residues and mutations are indicated. GAF: cyclic GMP, adenylyl cyclase, FhlA. (B) The ETR1 receptor signal output is facilitated by the docking of CTR1 to the histidine kinase (HK) domain, where the HK domain inhibits receptor signaling. (C) The deletion or mutation of the HK domain may result in a conformational change that enables ETR1 receptor signal output in a CTR1-independent manner to inhibit EIN2-activated ethylene signaling. (D) Without activation by CTR1, the ETR1 receptor is capable of a basal-level receptor signaling, partly alleviating EIN2 activity. (E) In the absence of the ethylene receptor, EIN2 is fully active. (F) The ethylene receptor is inactivated by the binding of excess CTR1-N, and EIN2 is relieved from inhibition to activate ethylene signaling. Arrowheads of different grayscales, from weak to strong, indicate the strength of EIN2 activity. In this model, ethylene receptor isoforms are represented by ETR1. With little information about the properties and identity of the EIN2 fragments, the fragments are not illustrated in this mode.

Ethylene signaling has been investigated for more than three decades under the guidance of the present genetic framework, and its biochemical nature has yet to be determined. In this study, we present experimental results that expand the knowledge of the present framework and propose a different model explaining how ethylene signaling is regulated by the ethylene receptor isoforms, CTR1, and EIN2. The underlying molecular events involved in ethylene signaling are more complicated than previously proposed, and our study may help future studies develop a niche that provides other insights into ethylene signaling.

## Acknowledgments

We thank Dr. J-S Zhang of CAS for the *mhl1*, *mhl2*, *mhl1 mhl2*, and *mhl1 mhl2 ctr1-1* germplasms (13). Ms. ZW Xiang was involved in part of the work on *ETP1/ETP2*. This work was supported by the National Science Foundation of China (NSFC; NSFC:32070315 and 32270325) and the Chinese Academy of Sciences (XDB27030208) to CK Wen. LSCM was conducted at the facility center of the CAS Center for Excellence in Molecular Plant Sciences.

## Author contributions

Designed the research and data analyses: C-K Wen and HW Zhao; Performed the research: HW Zhao, Y Zhang, Y Chen, C Wang, Q Liu, and JY Zhang; C-K Wen wrote the paper.

### Data Availability Statement

All data supporting the findings of this study are available within the paper and its Supplementary Information. Source data are provided with this paper.

---

**Supplementary Table S1.** Primer sets used in this study.

**Supplemental Figure S1.**
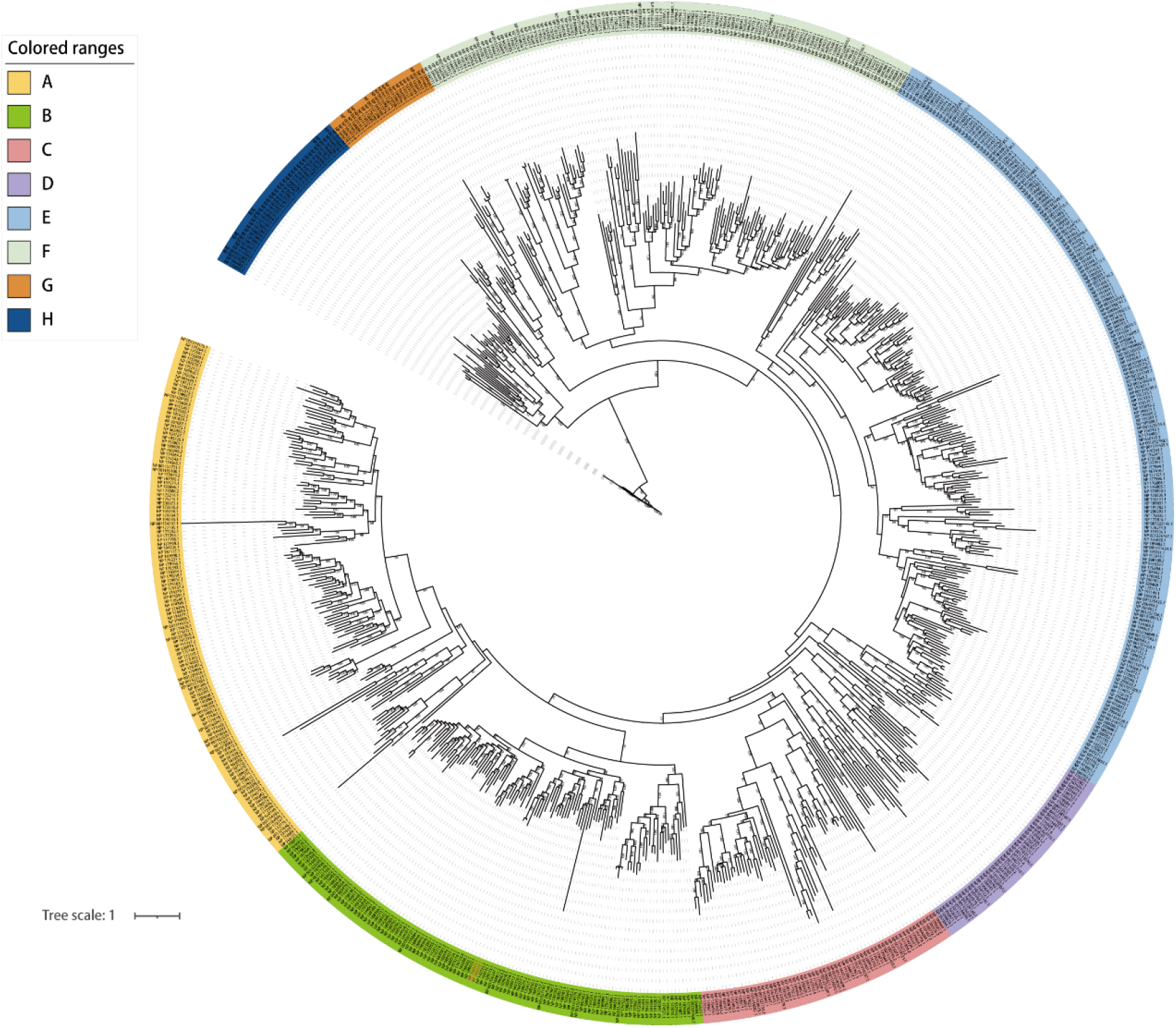
A phylogeny of the 745 F-box-containing proteins. The 745 F-box-containing proteins can be categorized into eight subclades, highlighted with different colors.

**Supplemental Figure S2.**
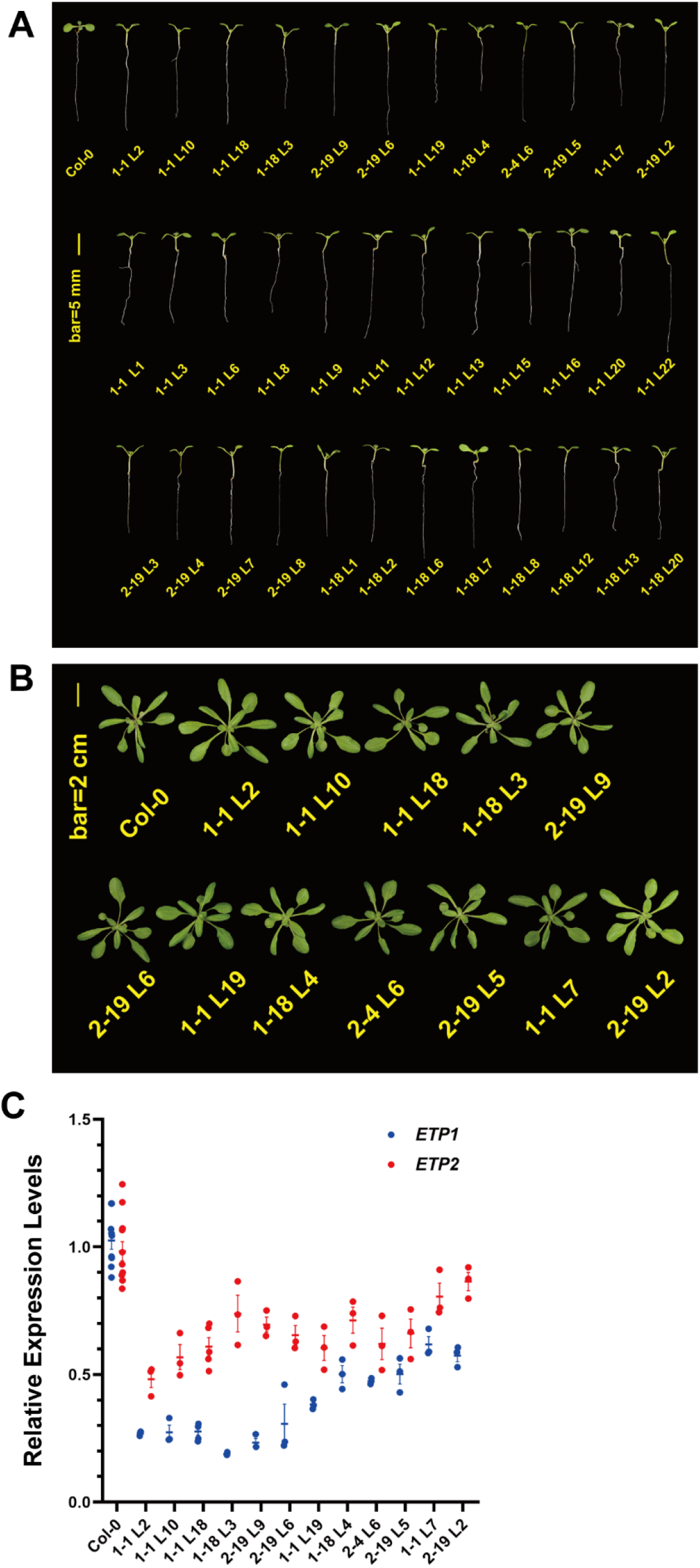
*amiRNA* overexpression has little impact on seedling growth. (A) The phenotypes of 36 independent *amiRNA*-overexpressing seedling lines. Rosette phenotype (B) and RT-qPCR measurements of *ETP1/ETP2* (C) of the 12 independent *amiRNA*-overexpressing lines.

**Supplemental Figure S3.**
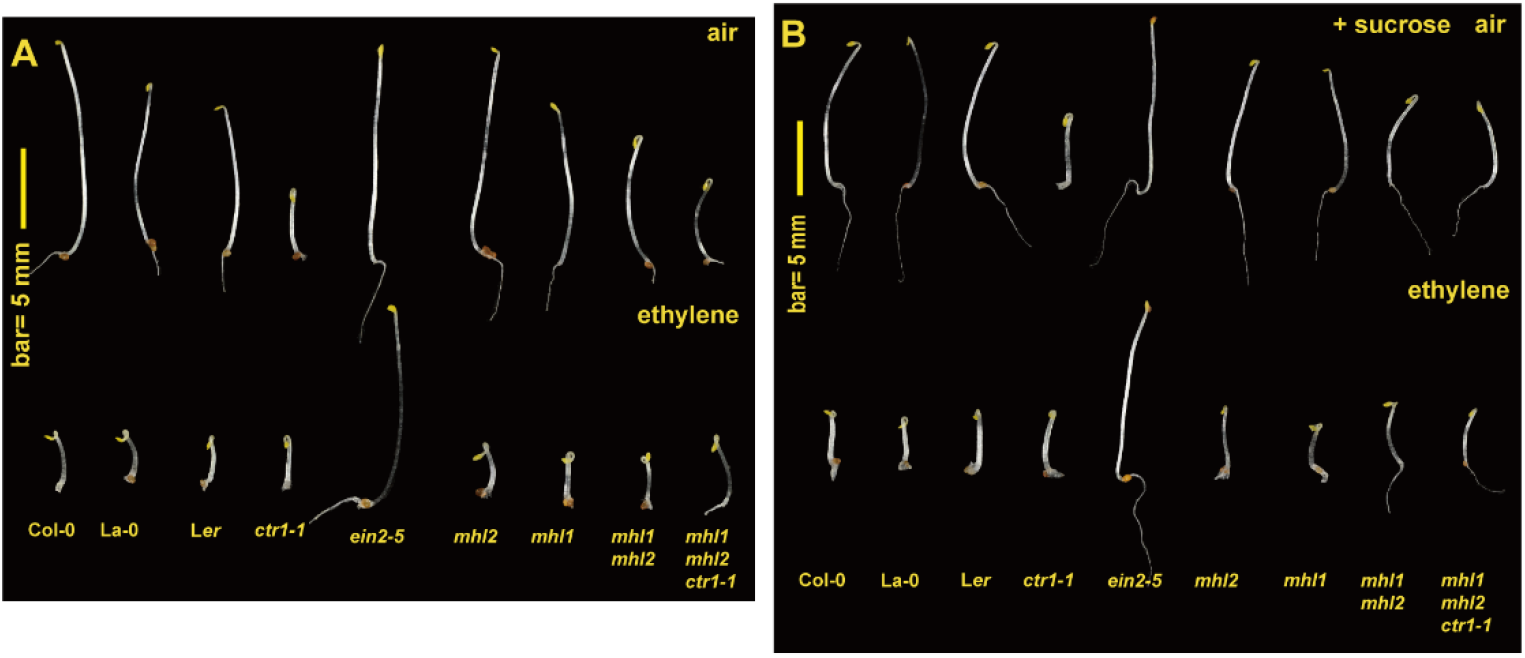
The *mhl1 mhl2* double mutation impacts seedling growth independent of ethylene. The phenotype of etiolated seedlings of the indicated genotypes, without (A) or with (B) sucrose supplementation, grown without (air) or with ethylene treatment.

**Supplemental Figure S4.**
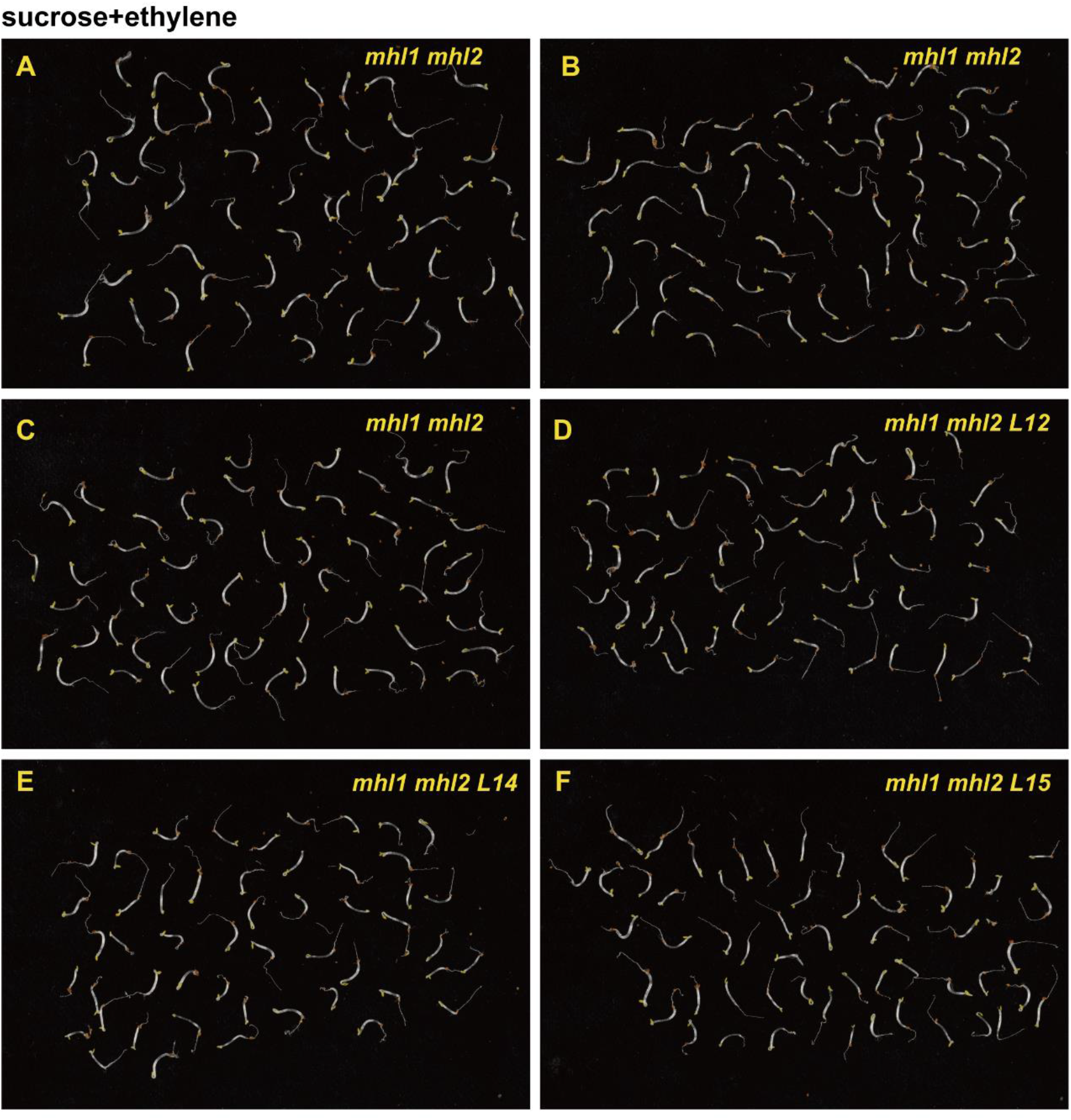
The root ethylene-insensitive phenotype is not associated with the *mhl1 mhl2* double mutation. (A)-(C) The seedling root phenotype of three independent progeny lines from the reported *mhl1 mhl2* germplasm (13). (D)-(F) The seedling root phenotype of three independent lines generated by the genetic crossing of *mhl1* and *mhl2*. Seedlings were grown in the dark with ethylene treatment and sucrose supplementation.

## Materials and Methods

### Plant materials and growth

Arabidopsis growth conditions and seedling hypocotyl measurements were performed as previously described (33). In brief, hypocotyl length was measured 80 hr after 72 hr of stratification for etiolated seedlings involving the use of Video tesT (Moscow) (21). The phenotypes of the light-grown seedlings and rosettes were scored at the 4-day and 4-week stages after germination, respectively. Arabidopsis protoplast preparation and protoplast transient expression were performed as previously described (49, 50). The *ctr1-1 ein2^W308*^/EIN2* and *ctr1-3 ein2^465*^/EIN2* sesquimutants were previously described (18), and the *etr1-10* mutant was previously reported (21). The *etr1-10 ers1^91*^* mutant was generated by genetic editing of the *etr1-10* mutant, with the transgene crossed out. Genetic editing of the *ERS2* gene in the *etr2-3 ein4-4* mutant generated the *etr2-3 ein4-4 ers2^146*^* mutant, and the transgene was subsequently crossed out. The receptor *ers1^91*^etr2-3 ein4-4 ers2^146*^* quadruple (*quad*) and quintuple (*quint*) mutants were isolated from the progeny of the genetic crosses of *etr1-10 ers1^91*^* and the *etr2-3 ein4-4 ers2^146*^* mutant. The *ctr1-3 quad* and *ctr1-3 qunit* mutants were isolated from the progeny of the genetic crossing of *ETR1^1-349^ ers1^91*^etr2-3 ein4-4 ers2^146*^*and the *etr1-10 ers1^91*^ ctr1-3* mutant. The *quint ein2^W308*^* mutant was generated from the genetic crossing of *etr1-10 ers1^91*^ ein2^W308*^* and *ers1^91*^etr2-3 ein4-4 ers2^146*^*. Among those mutants, the mutants with the *etr1-10 ers1^91*^*double mutation are lethal and infertile and kept as heterozygous.

### Clones and genetic editing

Primer information for RT‒qPCR, cloning, genotyping, and gene editing are listed in Supplemental Table S1. If not otherwise specified, ethylene treatment involved the addition of 10 µL L^-1^ ethylene. RT‒qPCR was performed with StepOne Plus (Applied Biosystems) and Takara SYBR Premix Ex Taq. The resulting clones were subsequently transformed with the binary vector pCAMBIA1300 or pCGN1547 in *Agrobacterium*. The clones in *Agrobacterium* were confirmed by retransforming to *E. coli*, and the clones were examined to confirm that the clones were not rearranged in *Agrobacterium*. Mutants obtained from CRISPR-Cas9-mediated editing were backcrossed with wild type (Col-0) to remove the transgenes. The resulting mutations were confirmed by sequencing. Among the several *etp1* and *etp2* alleles (Fig. 2A), the *etp1^7*^ etp2^41*-1^* double mutation (designated *etp1 etp2*) was investigated. For transgenes, the *UBQ10* promoter was involved for the *ETP1*, *ETP2*, *MHL1*, and *MHL2* transgenes, and the *EIN2* promoter was involved for the *EIN2* and *EIN2* variant transgenes, except for the *EIN2-eGFP* transgene, which involved the *SUPER* promoter. For transient expression assays in Arabidopsis protoplasts, the *35S* promoter of the pA7 vector was used. The *amiRNA* clone was constructed as previously described (2).

### Sequence analyses

The signal peptide, transmembrane domain, and sequence alignment matrix for MHL1, MHL2, and rice MHZ3 were determined from UniProt (https://www.uniprot.org/)(1). The reported *MHZ3* locus LOC_Os06g02480 (13) could not be retrieved from the NCBI GenBank, and the accession Os06g0115200 for *MHZ3* was identified. The accessions for MHZ3, MHL1, and MHL2 are Q8H668, Q9FRK5, and Q9LN61, respectively, at UniProt. The structures of ETP1, ETP2, and the ETP-related proteins were determined by UniProt. The possible targeting of the amiRNA to the 19 *ETP*-related transcripts was determined following previously described criteria (12).

#### Phylogenesis of Arabidopsis F-box-containing proteins

The protein sequence of the F-box gene family from *Arabidopsis thaliana* used was obtained as previously reported (34), with the redundant sequences removed, and a total of 745 full-length amino acid sequences were aligned by MAFFT (version 7.520) (51) involving the autooption strategy. The alignment was input into ModelFinder, a program implemented in IQtree (version 2.2.2.6)(52), to assess the best-fit substitution model according to the Bayesian information criterion (BIC), and the JTT+F+R9 substitution model was chosen to construct the maximum likelihood (ML) tree. Tree branch support was established through 1,000 iterations of UltraFast bootstraps. The resulting tree was further visualized and edited using the online program iTOL v6 (https://itol.embl.de/).

### Immunoassay and antibodies

For the immunoassay, total protein from crude extracts (50 mM Tris·HCl, pH 7.5, 150 mM NaCl, 5 mM EDTA·2Na, 0.1% Triton X-100, 0.2% NP-40, 1 mM PMSF, 1 mM cocktail, 1 mM DTT) of plant materials was mixed with 6× SDS buffer. Membrane protein enrichment involved pulverization of Arabidopsis seedlings (2-week stage) in liquid nitrogen, homogenization in extraction buffer, and filtration through a nylon membrane. The filtrate was centrifuged at 3,000×g at 4 °C for 20 min, and the resulting supernatant was centrifuged at 8,000×g at 4 °C for 15 min. The resulting supernatant was subjected to ultracentrifugation at 100,000×g at 4 °C for 1 hr with a TYPE 90 Ti rotor. The resulting supernatant was the cytoplasmic soluble fraction. The membrane proteins were sonicated and solubilized with 6× SDS buffer at 37 °C for 1 hr. To enrich the nuclear proteins, the homogenized cell extract collected after centrifugation at 8,000×g was resuspended in Buffer A (10 mM HEPES, pH 7.8; 10 mM KCl; 10 mM MgCl_2_; 5 mM EDTA; 250 mM sucrose; 0.5% Triton X-100; 1 mM DTT; and 0.2 mM PMSF), followed by centrifugation at 4 °C at 2,000× for 15 min. The pellet was resuspended in Buffer A, and this procedure was repeated 5 times. The resulting nuclear pellet was resuspended, sonicated, and centrifuged at 16,000×g at 4 °C for 5 min. The resulting supernatant was dissolved in 6× SDS buffer and solubilized at 37 °C for 1 hr. Details for protein transient expression in Arabidopsis protoplasts and coimmunoprecipitation were followed as described (18, 40).

For western blots, the protein was solubilized at 37 °C instead of at 100 °C to ensure better solubilization and prevent protein aggregation of the membrane protein (53). The EIN2 variant proteins were detected by polyclonal anti-GFP antibodies [rabbit anti-GFP tag pAb, ABclonal or mouse monoclonal anti-GFP (JL-8; Clontech, 632381)] with a LumiBest ECL substrate solution kit (Share-bio, Shanghai) as the substrate for horseradish peroxidase and goat anti-rabbit IgGs. Chemiluminescence was acquired by Tanon-5500Multi (Tanon, Shanghai). Details for protein isolation were described (18). Sucrose density gradient fractionation was performed with a Gradient Maker (Gradient Master 108, BioComp, Canada). Details for the enrichment of the membrane and nuclear fractions and the density gradient fractionation are described (40).

#### Laser scanning confocal microscopy (LSCM)

Laser scanning confocal microscopy (LSCM) was performed with a Zeiss LSM880 or Leica TSC SP8 STED 3× instrument provided by the Facility Center of CEMPS (CAS Center for Excellence in Molecular Plant Sciences). The eYFP/YFP fluorescence was acquired by 514 nm laser excitation, and the emission was recorded between 520 and 540 nm. The eGFP/GFP fluorescence was detected by 488 nm laser excitation, and the emission was recorded between 498 and 518 nm. mRFP/mCherry fluorescence was detected by 543 nm laser excitation, and the emission was recorded between 590 and 640 nm. Transgenic Arabidopsis cells from etiolated seedling hypocotyls were subjected to LSCM. Transient expression involved Arabidopsis protoplasts (18).

### Statistics

Statistical comparisons of multiple sample means involved one-way ANOVA and *a posteriori* multiple comparisons, and the mean differences are presented as 95% confidence intervals (CI_0.95_) for the interpretation of the biological significance or effect size (33, 54). The sample sizes, if not specified, were greater than 30 (*n*>30) for seedling hypocotyl measurements and ≥3 for RT‒qPCR from independent samples; each replicate included >50 seedlings. The statistical analyses and graphs were generated by GraphPad Prism 8.0.

